# Geometric models reveal behavioral and neural signatures of transforming naturalistic experiences into episodic memories

**DOI:** 10.1101/409987

**Authors:** Andrew C. Heusser, Paxton C. Fitzpatrick, Jeremy R. Manning

**Affiliations:** Department of Psychological and Brain Sciences, Dartmouth College, Hanover, NH 03755, USA; Akili Interactive Labs, Boston, MA 02110

## Abstract

The mental contexts in which we interpret experiences are often person-specific, even when the experiences themselves are shared. We developed a geometric framework for mathematically characterizing the subjective conceptual content of dynamic naturalistic experiences. We model experiences and memories as *trajectories* through word embedding spaces whose coordinates reflect the universe of thoughts under consideration. Memory encoding can then be modeled as geometrically preserving or distorting the *shape* of the original experience. We applied our approach to data collected as participants watched and verbally recounted a television episode while undergoing functional neuroimaging. Participants’ recountings all preserved coarse spatial properties (essential narrative elements), but not fine spatial scale (low-level) details, of the episode’s trajectory. We also identified networks of brain structures sensitive to these trajectory shapes. Our work provides insights into how we preserve and distort our ongoing experiences when we encode them into episodic memories.

## Introduction

What does it mean to *remember* something? In traditional episodic memory experiments (e.g., list-learning or trial-based experiments; Kahana, 1996; Murdock, 1962), remembering is often cast as a discrete, binary operation: each studied item may be separated from the rest of one’s experience and labeled as having been either recalled or forgotten. More nuanced studies might incorporate self-reported confidence measures as a proxy for memory strength, or ask participants to discriminate between recollecting the (contextual) details of an experience and having a general feeling of familiarity (Yonelinas, 2002). Using well-controlled, trial-based experimental designs, the field has amassed a wealth of information regarding human episodic memory (for review see Kahana, 2012). However, there are fundamental properties of the external world and our memories that trial-based experiments are not well suited to capture (for review, also see Huk et al., 2018; Koriat and Goldsmith, 1994). First, our experiences and memories are continuous, rather than discrete— isolating a naturalistic event from the context in which it occurs can substantially change its meaning. Second, whether or not the rememberer has precisely reproduced a specific set of words in describing a given experience is nearly orthogonal to how well they were actually able to remember it. In classic (e.g., list-learning) memory studies, by contrast, the number or proportion of *exact* recalls is often considered to be a primary metric for assessing the quality of participants’ memories. Third, one might remember the essence (or a general summary) of an experience but forget (or neglect to recount) particular low-level details. Capturing the essence of what happened is often a main goal of recounting an episodic memory to a listener, whereas the inclusion of specific low-level details is often less pertinent.

How might we formally characterize the *essence* of an experience, and whether it has been recovered by the rememberer? And how might we distinguish an experience’s overarching essence from its low-level details? One approach is to start by considering some fundamental properties of the dynamics of our experiences. Each given moment of an experience tends to derive meaning from surrounding moments, as well as from longer-range temporal associations (Lerner et al., 2011; Manning, 2019, 2020). Therefore, the timecourse describing how an event unfolds is fundamental to its overall meaning. Further, this hierarchy formed by our subjective experiences at different timescales defines a *context* for each new moment (e.g., Howard and Kahana, 2002; Howard et al., 2014), and plays an important role in how we interpret that moment and remember it later (for review see Manning, 2020; Manning et al., 2015). Our memory systems can leverage these associations to form predictions that help guide our behaviors (Ranganath and Ritchey, 2012). For example, as we navigate the world, the features of our subjective experiences tend to change gradually (e.g., the room or situation we find ourselves in at any given moment is strongly temporally autocorrelated), allowing us to form stable estimates of our current situation and behave accordingly (Zacks et al., 2007; Zwaan and Radvansky, 1998).

Occasionally, this gradual drift of our ongoing experience is punctuated by sudden changes, or shifts (e.g., when we walk through a doorway; Radvansky and Zacks, 2017). Prior research suggests that these sharp transitions (termed *event boundaries*) help to discretize our experiences (and their mental representations) into *events* (Brunec et al., 2018; Clewett and Davachi, 2017; DuBrow and Davachi, 2013; Ezzyat and Davachi, 2011; Heusser et al., 2018a; Radvansky and Zacks, 2017). The interplay between the stable (within-event) and transient (across-event) temporal dynamics of an experience also provides a potential framework for transforming experiences into memories that distills those experiences down to their essences. For example, prior work has shown that event boundaries can influence how we learn sequences of items (DuBrow and Davachi, 2013; Heusser et al., 2018a), navigate (Brunec et al., 2018), and remember and understand narratives (Ezzyat and Davachi, 2011; Zwaan and Radvansky, 1998). This work also suggests a means of distinguishing the essence of an experience from its low-level details: The overall structure of events and event transitions reflects how the high-level experience unfolds (i.e., its essence), while subtler event-level properties reflect its low-level details. Prior research has also implicated a network of brain regions (including the hippocampus and the medial prefrontal cortex) in playing a critical role in transforming experiences into structured and consolidated memories (Tompary and Davachi, 2017).

Here, we sought to examine how the temporal dynamics of a naturalistic experience were later reflected in participants’ memories. We also sought to leverage the above conceptual insights into the distinctions between an experience’s essence and its low-level details to build models that explicitly quantified these distinctions. We analyzed an open dataset that comprised behavioral and functional Magnetic Resonance Imaging (fMRI) data collected as participants viewed and then verbally recounted an episode of the BBC television show Sherlock (Chen et al., 2017). We developed a computational framework for characterizing the temporal dynamics of the moment-by-moment content of the episode and of participants’ verbal recalls. Our framework uses topic modeling (Blei et al., 2003) to characterize the thematic conceptual (semantic) content present in each moment of the episode and recalls by projecting each moment into a word embedding space. We then use hidden Markov models (Baldassano et al., 2017; Rabiner, 1989) to discretize this evolving semantic content into events. In this way, we cast both naturalistic experiences and memories of those experiences as geometric *trajectories* through word embedding space that describe how they evolve over time. Under this framework, successful remembering entails verbally traversing the content trajectory of the episode, thereby reproducing the shape (essence) of the original experience. Our framework captures the episode’s essence in the sequence of geometric coordinates for its events, and its low-level details by examining its within-event geometric properties.

Comparing the overall shapes of the topic trajectories for the episode and participants’ recalls reveals which aspects of the episode’s essence were preserved (or lost) in the translation into memory. We also develop two metrics for assessing participants’ memories for low-level details: (1) the *precision* with which a participant recounts details about each event, and (2) the *distinctiveness* of their recall for each event, relative to other events. We examine how these metrics relate to overall memory performance as judged by third-party human annotators. We also compare and contrast our general approach to studying memory for naturalistic experiences with standard metrics for assessing performance on more traditional memory tasks, such as list-learning. Last, we leverage our framework to identify networks of brain structures whose responses (as participants watched the episode) reflected the temporal dynamics of the episode and/or how participants would later recount it.

## Results

To characterize the dynamic content of the *Sherlock* episode and participants’ subsequent recountings, we used a topic model (Blei et al., 2003) to discover the episode’s latent themes. Topic models take as inputs a vocabulary of words to consider and a collection of text documents, and return two output matrices. The first of these is a *topics matrix* whose rows are *topics* (or latent themes) and whose columns correspond to words in the vocabulary. The entries in the topics matrix reflect how each word in the vocabulary is weighted by each discovered topic. For example, a detective-themed topic might weight heavily on words like “crime,” and “search.” The second output is a *topic proportions matrix* with one row per document and one column per topic. The topic proportions matrix describes the mixture of discovered topics reflected in each document.

Chen et al. (2017) collected hand-annotated information about each of 1,000 (manually delineated) time segments spanning the roughly 50 minute video used in their study. Each annotation included: a brief narrative description of what was happening, the location where the action took place, the names of any characters on the screen, and other similar details (for a full list of annotated features, see *Methods*). We took the union of all unique words (excluding stop words, such as “and,” “or,” “but,” etc.) across all features from all annotations as the vocabulary for the topic model. We then concatenated the sets of words across all features contained in overlapping sliding windows of (up to) 50 annotations, and treated each window as a single document for the purpose of fitting the topic model. Next, we fit a topic model with (up to) *K* = 100 topics to this collection of documents. We found that 32 unique topics (with non-zero weights) were sufficient to describe the time-varying content of the episode (see *Methods*; Figs. 1, S2). We note that our approach is similar in some respects to Dynamic Topic Models (Blei and Lafferty, 2006) in that we sought to characterize how the thematic content of the episode evolved over time. However, whereas Dynamic Topic Models are designed to characterize how the properties of *collections* of documents change over time, our sliding window approach allows us to examine the topic dynamics within a single document (or video). Specifically, our approach yielded (via the topic proportions matrix) a single *topic vector* for each sliding window of annotations transformed by the topic model. We then stretched (interpolated) the resulting windows-by-topics matrix to match the time series of the 1,976 fMRI volumes collected as participants viewed the episode.

**Figure 1:**
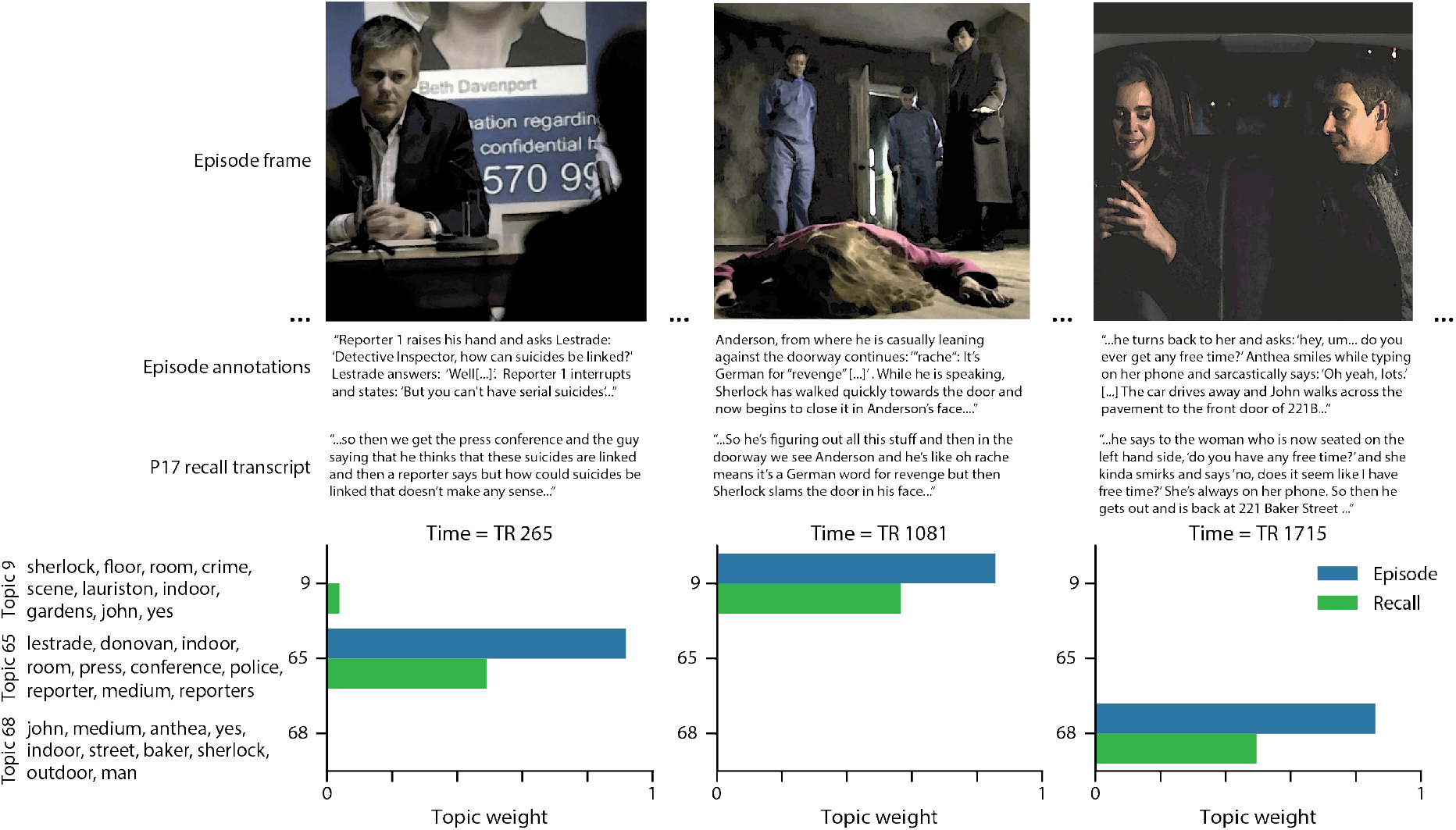
Topic weights in episode and recall content. We used detailed, hand-generated annotations describing each manually identified time segment from the episode to fit a topic model. Three example frames from the episode (first row) are displayed, along with their descriptions from the corresponding episode annotation (second row) and an example participant’s recall transcript (third row). We used the topic model (fit to the episode annotations) to estimate topic vectors for each moment of the episode and each sentence of participants’ recalls. Example topic vectors are displayed in the bottom row (blue: episode annotations; green: example participant’s recalls). Three topic dimensions are shown (the highest-weighted topics for each of the three example scenes, respectively), along with the 10 highest-weighted words for each topic. Figure S2 provides a full list of the top 10 words from each of the discovered topics.

The 32 topics we found were heavily character-focused (i.e., the top-weighted word in each topic was nearly always a character) and could be roughly divided into themes centered around Sherlock Holmes (the titular character), John Watson (Sherlock’s close confidant and assistant), supporting characters (e.g., Inspector Lestrade, Sergeant Donovan, or Sherlock’s brother Mycroft), or the interactions between various groupings of these characters (Fig. S2). This likely follows from the frequency with which these terms appeared in the episode annotations. Several of the identified topics were highly similar, which we hypothesized might allow us to distinguish between subtle narrative differences if the distinctions between those overlapping topics were meaningful. The topic vectors for each timepoint were also *sparse*, in that only a small number of topics (typically one or two) tended to be “active” in any given timepoint (Fig. 2A). Further, the dynamics of the topic activations appeared to exhibit *persistence* (i.e., given that a topic was active in one timepoint, it was likely to be active in the following timepoint) along with *occasional rapid changes* (i.e., occasionally topic weights would change abruptly from one timepoint to the next). These two properties of the topic dynamics may be seen in the block diagonal structure of the timepoint-by-timepoint correlation matrix (Fig. 2B) and reflect the gradual drift and sudden shifts fundamental to the temporal dynamics of many real-world experiences, as well as television episodes. Given this observation, we adapted an approach devised by Baldassano et al. (2017), and used a hidden Markov model (HMM) to identify the *event boundaries* where the topic activations changed rapidly (i.e., the boundaries of the blocks in the temporal correlation matrix; event boundaries identified by the HMM are outlined in yellow in Fig. 2B). Part of our model fitting procedure required selecting an appropriate number of events into which the topic trajectory should be segmented. To accomplish this, we used an optimization procedure that maximized the difference between the topic weights for timepoints within an event versus timepoints across multiple events (see *Methods*). We then created a stable summary of the content within each episode event by averaging the topic vectors across the timepoints spanned by each event (Fig. 2C).

**Figure 2:**
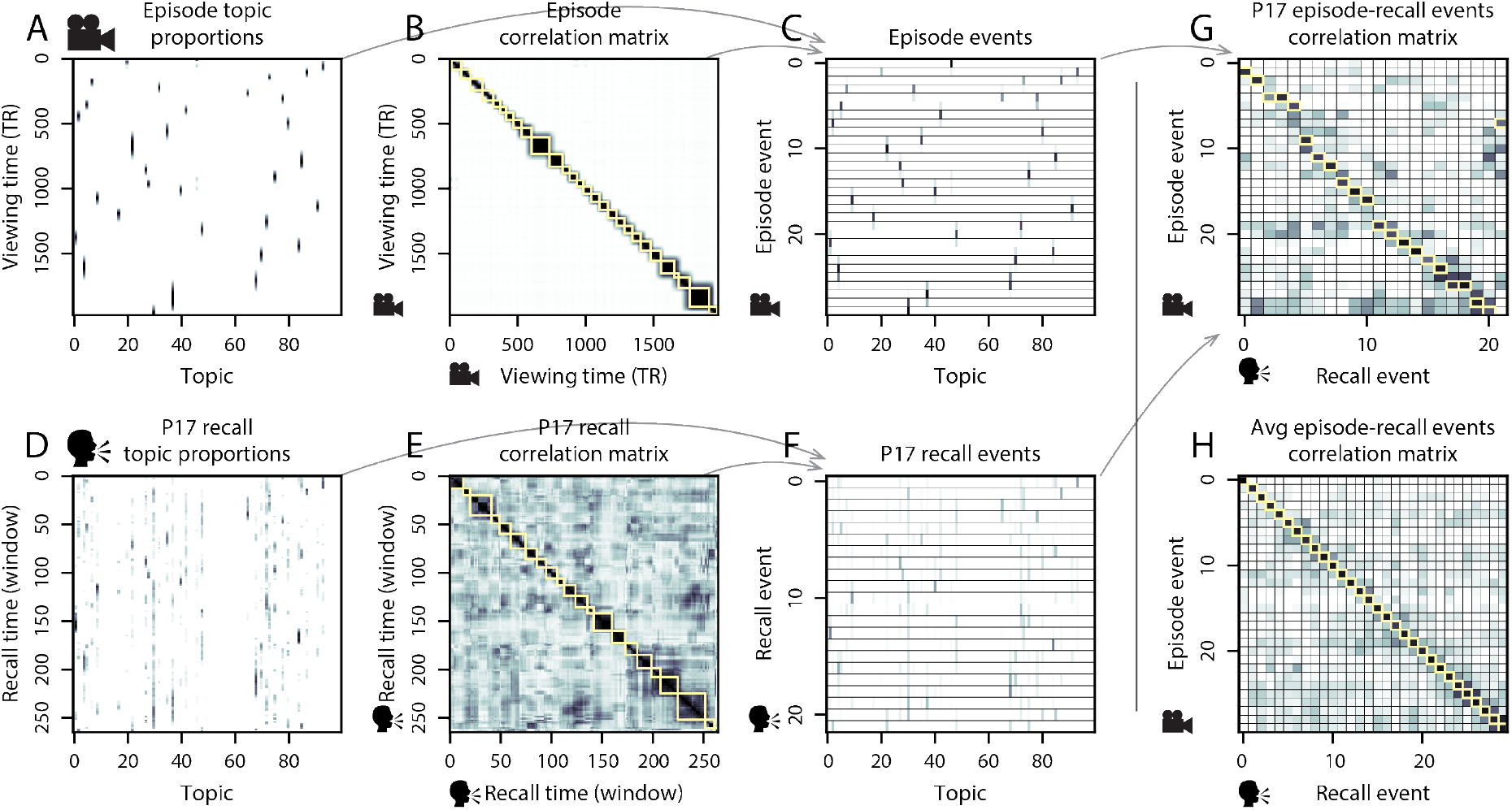
Modeling naturalistic stimuli and recalls. All panels: darker colors indicate greater values; range: [0, 1]. **A**. Topic vectors (*K* = 100) for each of the 1976 episode timepoints. **B**. Timepoint-by-timepoint correlation matrix of the topic vectors displayed in Panel A. Event boundaries discovered by the HMM are denoted in yellow (30 events detected). **C**. Average topic vectors for each of the 30 episode events. **D**. Topic vectors for each of 265 sliding windows of sentences spoken by an example participant while recalling the episode. **E**. Timepoint-by-timepoint correlation matrix of the topic vectors displayed in Panel D. Event boundaries detected by the HMM are denoted in yellow (22 events detected). For similar plots for all participants, see Figure S4. **F**. Average topic vectors for each of the 22 recall events from the example participant. **G**. Correlations between the topic vectors for every pair of episode events (Panel C) and recall events (from the example participant; Panel F). For similar plots for all participants, see Figure S5. **H**. Average correlations between each pair of episode events and recall events (across all 17 participants). To create the figure, each recalled event was assigned to the episode event with the most correlated topic vector (yellow boxes in panels G and H).

Given that the time-varying content of the episode could be segmented cleanly into discrete events, we wondered whether participants’ recalls of the episode also displayed a similar structure. We applied the same topic model (already trained on the episode annotations) to each participant’s recalls. Analogously to how we parsed the time-varying content of the episode, to obtain similar estimates for each participant’s recall transcript, we treated each overlapping window of (up to) 10 sentences from their transcript as a document, and computed the most probable mix of topics reflected in each timepoint’s sentences. This yielded, for each participant, a number-of-windows by number-of-topics topic proportions matrix that characterized how the topics identified in the original episode were reflected in the participant’s recalls. An important feature of our approach is that it allows us to compare participants’ recalls to events from the original episode, despite that different participants used widely varying language to describe the events, and that those descriptions often diverged in content, quality, and quantity from the episode annotations. This ability to match up conceptually related text that differs in specific vocabulary, detail, and length is an important benefit of projecting the episode and recalls into a shared topic space. An example topic proportions matrix from one participant’s recalls is shown in Figure 2D.

Although the example participant’s recall topic proportions matrix has some visual similarity to the episode topic proportions matrix, the time-varying topic proportions for the example participant’s recalls are not as sparse as those for the episode (compare Figs. 2A and D). Similarly, although there do appear to be periods of stability in the recall topic dynamics (i.e., most topics are active or inactive over contiguous blocks of time), the changes in topic activations that define event boundaries appear less clearly delineated in participants’ recalls than in the episode’s annotations. To examine these patterns in detail, we computed the timepoint-by-timepoint correlation matrix for the example participant’s recall topic proportions matrix (Fig. 2E). As in the episode correlation matrix (Fig. 2B), the example participant’s recall correlation matrix has a strong block diagonal structure, indicating that their recalls are discretized into separated events. We used the same HMM-based optimization procedure that we had applied to the episode’s topic proportions matrix (see *Methods*) to estimate an analogous set of event boundaries in the participant’s recounting of the episode (outlined in yellow). We carried out this analysis on all 17 participants’ recall topic proportions matrices (Fig. S4).

Two clear patterns emerged from this set of analyses. First, although every individual participant’s recalls could be segmented into discrete events (i.e., every individual participant’s recall correlation matrix exhibited clear block diagonal structure; Fig. S4), each participant appeared to have a unique *recall resolution*, reflected in the sizes of those blocks. While some participants’ recall topic proportions segmented into just a few events (e.g., Participants P4, P5, and P7), others’ segmented into many shorter-duration events (e.g., Participants P12, P13, and P17). This suggests that different participants may be recalling the episode with different levels of detail—i.e., some might recount only high-level essential plot details, whereas others might recount low-level details instead (or in addition). The second clear pattern present in every individual participant’s recall correlation matrix was that, unlike in the episode correlation matrix, there were substantial off-diagonal correlations. Whereas each event in the original episode was (largely) separable from the others (Fig. 2B), in transforming those separable events into memory, participants appeared to be integrating across multiple events, blending elements of previously recalled and not-yet-recalled content into each newly recalled event (Figs. 2E, S4; also see Howard et al., 2012; Manning, 2019; Manning et al., 2011).

The above results demonstrate that topic models capture the dynamic conceptual content of the episode and participants’ recalls of the episode. Further, the episode and recalls exhibit event boundaries that can be identified automatically using HMMs to segment the dynamic content. Next, we asked whether some correspondence might be made between the specific content of the events the participants experienced while viewing the episode, and the events they later recalled. We labeled each recall event as matching the episode event with the most similar (i.e., most highly correlated) topic vector (Figs. 2G, S5). This yielded a sequence of “presented” events from the original episode, and a (potentially differently ordered) sequence of “recalled” events for each participant. Analogous to classic list-learning studies, we can then examine participants’ recall sequences by asking which events they tended to recall first (probability of first recall; Fig. 3A; Atkinson and Shiffrin, 1968; Postman and Phillips, 1965; Welch and Burnett, 1924); how participants most often transitioned between recalls of the events as a function of the temporal distance between them (lag-conditional response probability; Fig. 3B; Kahana, 1996); and which events they were likely to remember overall (serial position recall analyses; Fig. 3C; Murdock, 1962). Some of the patterns we observed appeared to be similar to classic effects from the list-learning literature. For example, participants had a higher probability of initiating recall with early events (Fig. 3A) and a higher probability of transitioning to neighboring events with an asymmetric forward bias (Fig. 3B). However, unlike what is typically observed in list-learning studies, we did not observe patterns comparable to the primacy or recency serial position effects (Fig. 3C). We hypothesized that participants might be leveraging meaningful narrative associations and references over long timescales throughout the episode.

**Figure 3:**
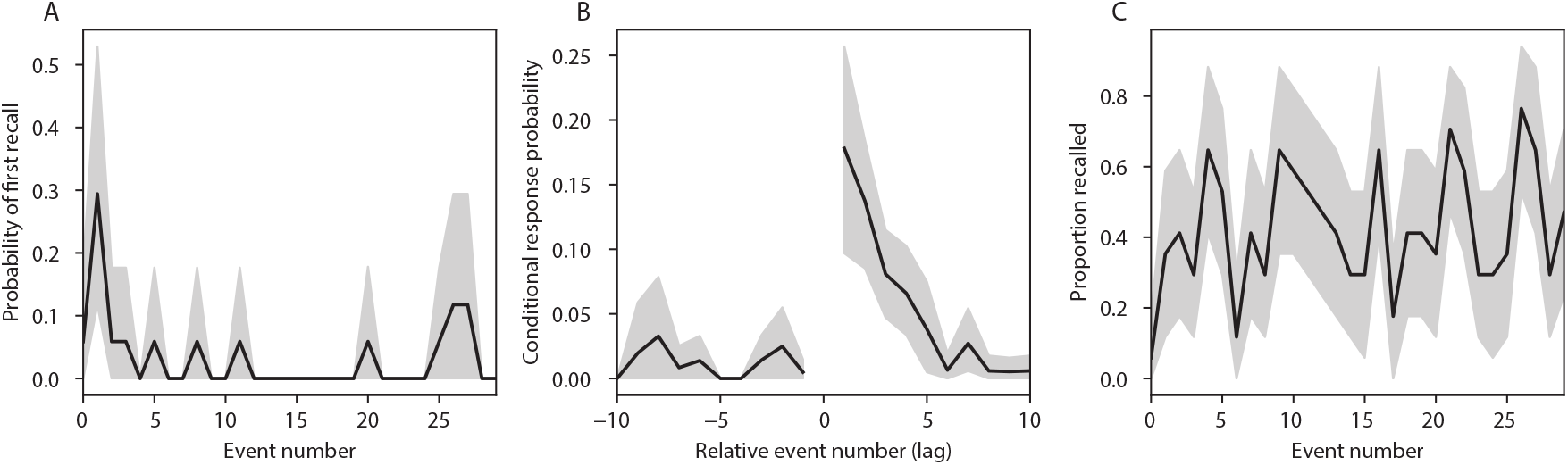
Naturalistic extensions of classic list-learning memory analyses. **A**. The probability of first recall as a function of the serial position of the event in the episode. **B**. The probability of recalling each event, conditioned on having most recently recalled the event *lag* events away in the episode. **C**. The proportion of participants who recalled each event, as a function of the serial position of the events in the episode. All panels: error ribbons denote bootstrap-estimated standard error of the mean.

Clustering scores are often used by memory researchers to characterize how people organize their memories of words on a studied list (for review, see Polyn et al., 2009). We defined analogous measures to characterize how participants organized their memories for episodic events (see *Methods* for details). Temporal clustering refers to the extent to which participants group their recall responses according to encoding position. Overall, we found that sequentially viewed episode events tended to appear nearby in participants’ recall event sequences (mean clustering score: 0.732, SEM: 0.033). Participants with higher temporal clustering scores tended to exhibit better overall memory for the episode, according to both Chen et al. (2017)’s hand-counted numbers of recalled scenes from the episode (Pearson’s *r*(15) = 0.49, *p* = 0.046) and the numbers of episode events that best-matched at least one recall event (i.e., model-estimated number of events recalled; Pearson’s *r*(15) = 0.59, *p* = 0.013). Semantic clustering measures the extent to which participants cluster their recall responses according to semantic similarity. We found that participants tended to recall semantically similar episode events together (mean clustering score: 0.650, SEM: 0.032), and that semantic clustering score was also related to both hand-counted (Pearson’s *r*(15) = 0.65, *p* = 0.005) and model-estimated (Pearson’s *r*(15) = 0.58, *p* = 0.015) numbers of recalled events.

The above analyses illustrate how our framework for characterizing the dynamic conceptual content of naturalistic episodes enables us to carry out analyses that have traditionally been applied to much simpler list-learning paradigms. However, perhaps the most interesting aspects of memory for naturalistic episodes are those that have no list-learning analogs. The nuances of how one’s memory for an event might capture some details, yet distort or neglect others, is central to how we use our memory systems in daily life. Yet when researchers study memory in highly simplified paradigms, those nuances are not typically observable. We next developed two novel, continuous metrics, termed *precision* and *distinctiveness*, aimed at characterizing distortions in the conceptual content of individual recall events, and the conceptual overlap between how people described different events.

*Precision* is intended to capture the “completeness” of recall, or how fully the presented content was recapitulated in a participant’s recounting. We define a recall event’s precision as the maximum correlation between the topic proportions of that recall event and any episode event (Fig. 4). In other words, given that a recall event best matches a particular episode event, more precisely recalled events overlap more strongly with the conceptual content of the original episode event. When a given event is assigned a blend of several topics, as is often the case (Fig. 2), a high precision score requires recapitulating the relative topic proportions during recall.

**Figure 4:**
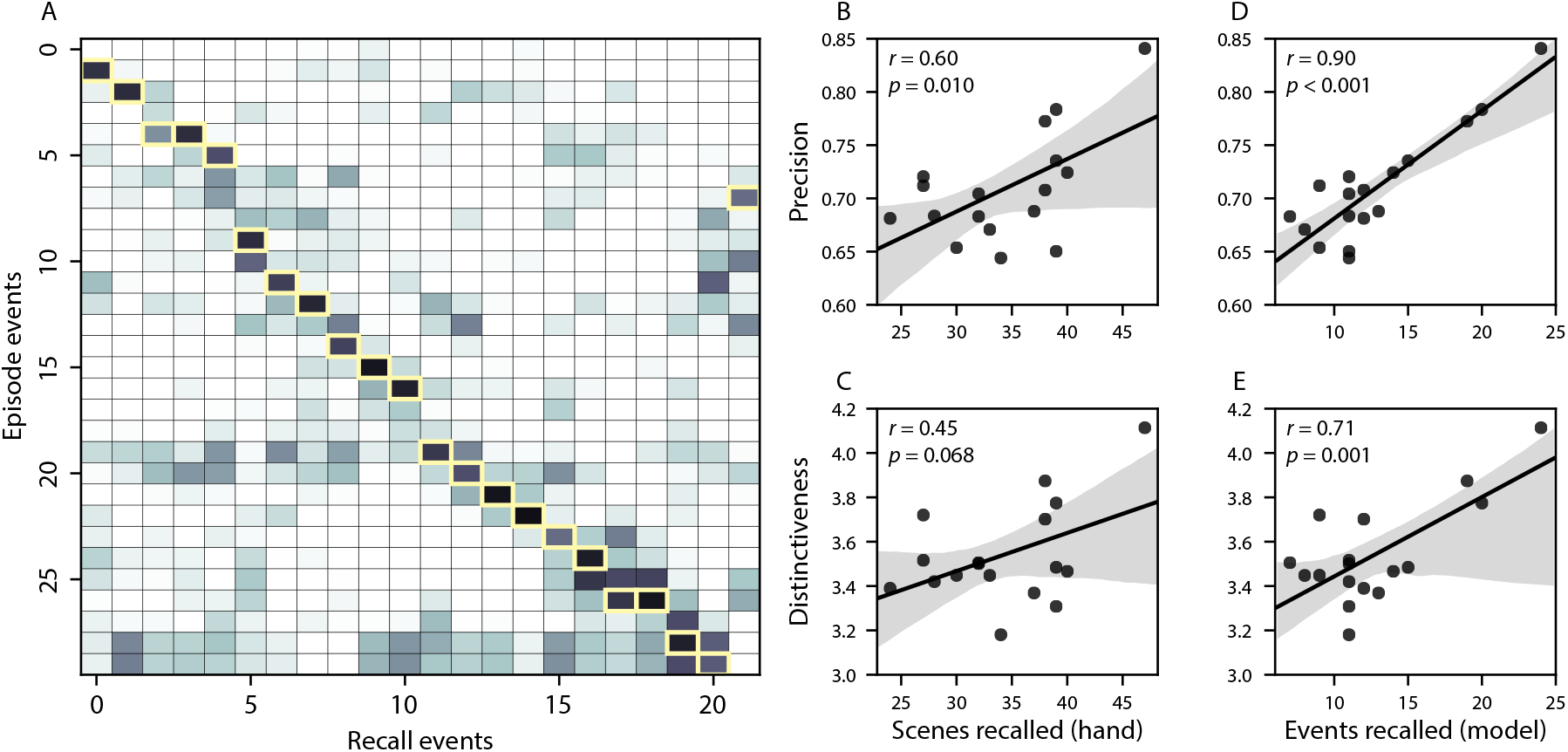
Novel content-based metrics of naturalistic memory: precision and distinctiveness. **A**. The episode-recall correlation matrix for a representative participant (P17). The yellow boxes highlight the maximum correlation in each column. The example participant’s overall precision score was computed as the average across the (Fisher *z*-transformed) correlation values in the yellow boxes. Their distinctiveness score was computed as the average (over recall events) of the *z*-scored (within column) event precisions. **B**. The (Pearson’s) correlation between precision and hand-counted number of recalled scenes. **C**. The correlation between distinctiveness and hand-counted number of recalled scenes. **D**. The correlation between precision and the number of recalled episode events, as determined by our model. **E**. The correlation between distinctiveness and the number of recalled episode events, as determined by our model.

*Distinctiveness* is intended to capture the “specificity” of recall. In other words, distinctiveness quantifies the extent to which a given recall event reflects the most similar episode event over and above other episode events. Intuitively, distinctiveness is like a normalized variant of our precision metric. Whereas precision solely measures how much detail about an episode was captured in someone’s recall, distinctiveness penalizes details that also pertain to other episode events. We define the distinctiveness of an event’s recall as its precision expressed in standard deviation units with respect to other episode events. Specifically, for a given recall event, we compute the correlation between its topic vector and that of each episode event. This yields a distribution of correlation coefficients (one per episode event). We subtract the mean and divide by the standard deviation of this distribution to *z*-score the coefficients. The maximum value in this distribution (which, by definition, belongs to the episode event that best matches the given recall event) is that recall event’s distinctiveness score. In this way, recall events that match one episode event far better than all other episode events will receive a high distinctiveness score. By contrast, a recall event that matches all episode events roughly equally will receive a comparatively low distinctiveness score.

In addition to examining how precisely and distinctively participants recalled individual events, one may also use these metrics to summarize each participant’s performance by averaging across a participant’s event-wise precision or distinctiveness scores. This enables us to quantify how precisely a participant tended to recall subtle within-event details, as well as how specific (distinctive) those details were to individual events from the episode. Participants’ average precision and distinctiveness scores were strongly correlated (*r*(15) = 0.90, *p* < 0.001). This indicates that participants who tended to precisely recount low-level details of episode events also tended to do so in an event-specific way (e.g., as opposed to detailing recurring themes that were present in most or all episode events; this behavior would have resulted in high precision but low distinctiveness). We found that, across participants, higher precision scores were positively correlated with the numbers of both hand-annotated scenes (*r*(15) = 0.60, *p* = 0.010) and model-estimated events (*r*(15) = 0.90, *p* < 0.001) that participants recalled. Participants’ average distinctiveness scores were also correlated with both the hand-annotated (*r*(15) = 0.45, *p* = 0.068) and model-estimated (*r*(15) = 0.71, *p* = 0.001) numbers of recalled events.

Examining individual recalls of the same episode event can provide insights into how the above precision and distinctiveness scores may be used to characterize similarities and differences in how different people describe the same shared experience. In Figure 5, we compare recalls for the same episode event from the participants with the highest (P17) and lowest (P6) precision scores. From the HMM-identified episode event boundaries, we recovered the set of annotations describing the content of a single episode event (event 21; Fig. 5C), and divided them into different color-coded sections for each action or feature described. Next, we used an analogous approach to identify the set of sentences comprising the corresponding recall event from each of the two example participants (Fig. 5D). We then colored all words describing actions and features in the transcripts shown in Panel D according to the color-coded annotations in Panel C. Visual comparison of these example recalls reveals that the more precise recall captures more of the episode event’s content, and in greater detail.

**Figure 5:**
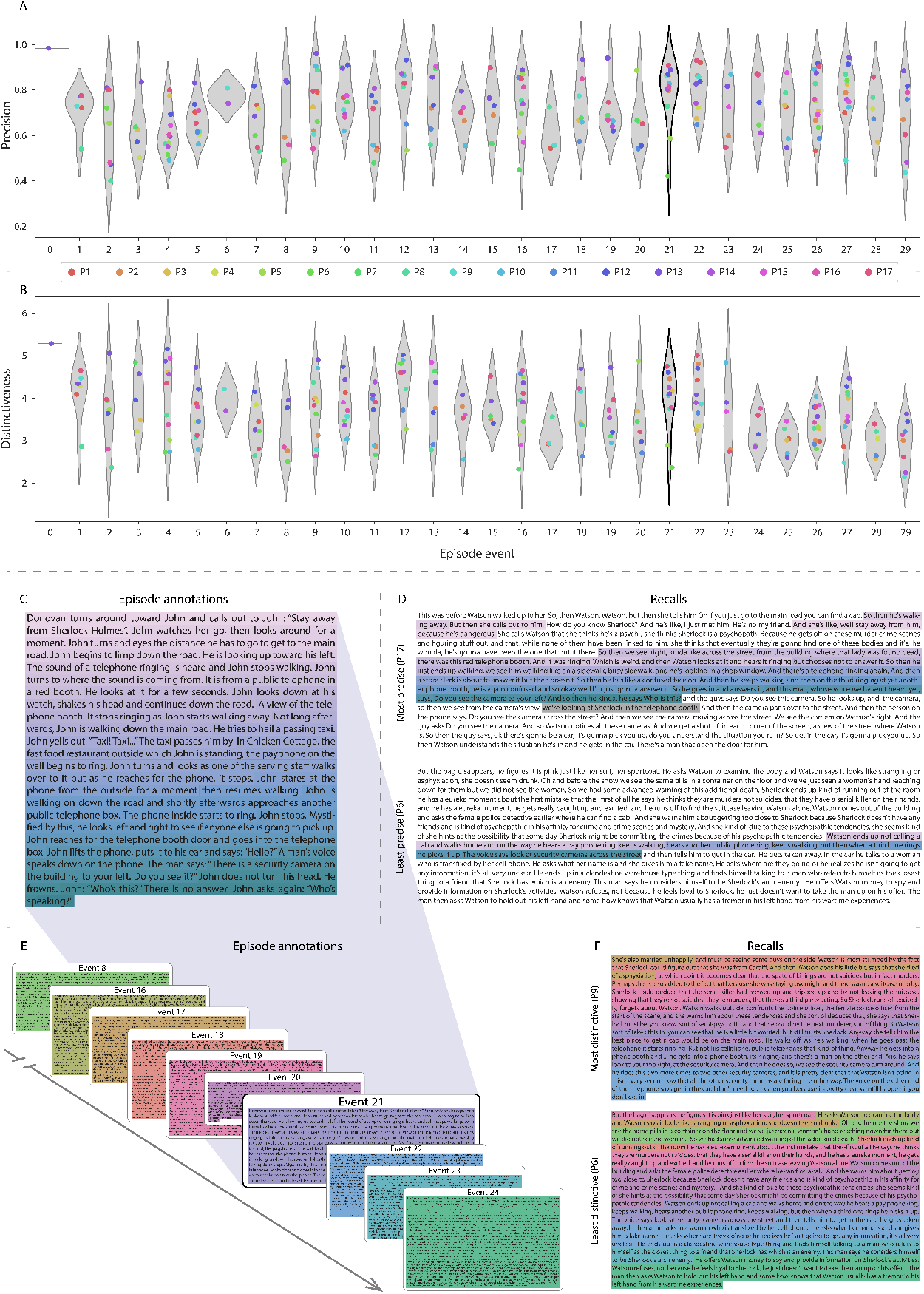
Precision reflects the completeness of recall, whereas distinctiveness reflects recall specificity. **A**. Recall precision by episode event. Grey violin plots display kernel density estimates for the distribution of recall precision scores for a single episode event. Colored dots within each violin plot represent individual participants’ recall precision for the given event. **B**. Recall distinctiveness by episode event, analogous to Panel A. **C**. The set of “Narrative Details” episode annotations (generated by Chen et al., 2017) comprising an example episode event (22) identified by the HMM. Each action or feature is highlighted in a different color. **D**. Sentences comprising the most precise (P17) and least precise (P6) participants’ recalls of episode event 21. Descriptions of specific actions or features reflecting those highlighted in Panel B are highlighted in the corresponding color. The text highlighted in gray denotes a (rare) false recall. The unhighlighted text denotes correctly recalled information about other episode events. **E**. The sets of “Narrative Details” episode annotations (generated by Chen et al., 2017) for scenes comprising episode events described by the example participants in Panel F. Each event’s text is highlighted in a different color. **F**. The sentences comprising the most distinctive (P9) and least distinctive (P6) participants’ recalls of episode event 21. Sections of recall describing each each episode event in Panel E are highlighted with the corresponding color.

Figure 5 also illustrates the differences between high and low distinctiveness scores. We extracted the set of sentences comprising the most distinctive recall event (P9) and least distinctive recall event (P6) corresponding to the example episode event shown in Panel C (event 21). We also extracted the annotations for all episode events whose content these participants’ single recall events described. We assigned each episode event a unique color (Fig. 5E), and colored each recalled sentence (Panel F) according to the episode events they best matched. Visual inspection of Panel F reveals that the most distinctive recall’s content is tightly concentrated around event 21, whereas the least distinctive recall incorporates content from a much wider range of episode events.

The preceding analyses sought to characterize how participants’ recountings of individual episode events captured the low-level details of each event. Next, we sought to characterize how participants’ recountings of the full episode captured its high-level essence—i.e., the shape of the episode’s trajectory through word embedding (topic) space. To visualize the essence of the episode and each participant’s recall trajectory (Heusser et al., 2018b), we projected the topic proportions matrices for the episode and recalls onto a shared two-dimensional space using Uniform Manifold Approximation and Projection (UMAP; McInnes et al., 2018). In this lower-dimensional space, each point represents a single episode or recall event, and the distances between the points reflect the distances between the events’ associated topic vectors (Fig. 6). In other words, events that are nearer to each other in this space are more semantically similar, and those that are farther apart are less so.

**Figure 6:**
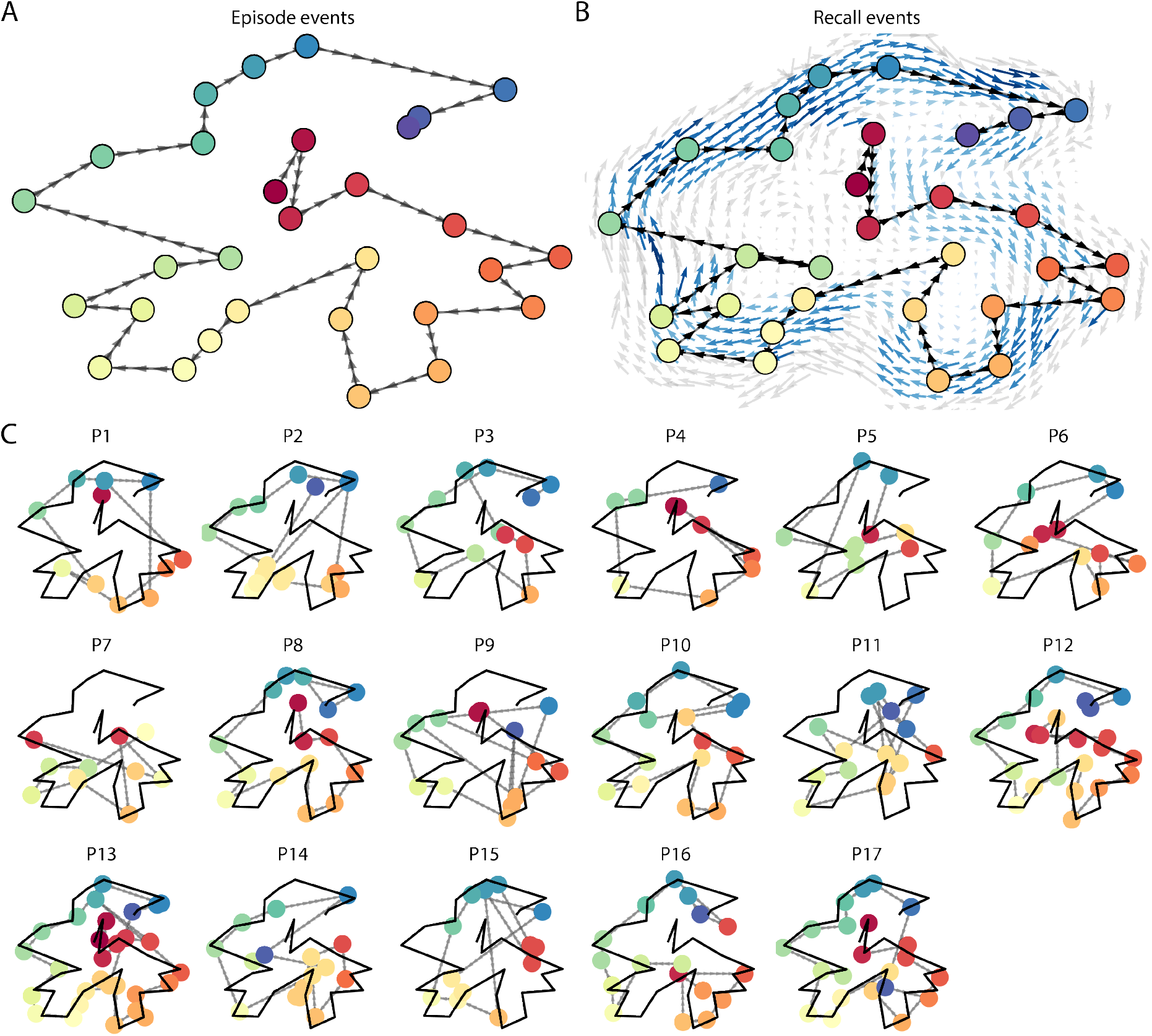
Trajectories through topic space capture the dynamic content of the episode and recalls. All panels: the topic proportion matrices have been projected onto a shared two-dimensional space using UMAP. **A**. The two-dimensional topic trajectory taken by the episode of *Sherlock*. Each dot indicates an event identified using the HMM (see *Methods*); the dot colors denote the order of the events (early events are in red; later events are in blue), and the connecting lines indicate the transitions between successive events. **B**. The average two-dimensional trajectory captured by participants’ recall sequences, with the same format and coloring as the trajectory in Panel A. To compute the event positions, we matched each recalled event with an event from the original episode (see *Results*), and then we averaged the positions of all events with the same label. The arrows reflect the average transition direction through topic space taken by any participants whose trajectories crossed that part of topic space; blue denotes reliable agreement across participants via a Rayleigh test (*p* < 0.05, corrected). **C**. The recall topic trajectories (gray) taken by each individual participant (P1–P17). The episode’s trajectory is shown in black for reference. Here, events (dots) are colored by their matched episode event (Panel A).

Visual inspection of the episode and recall topic trajectories reveals a striking pattern. First, the topic trajectory of the episode (which reflects its dynamic content; Fig. 6A) is captured nearly perfectly by the averaged topic trajectories of participants’ recalls (Fig. 6B). To assess the consistency of these recall trajectories across participants, we asked: given that a participant’s recall trajectory had entered a particular location in the reduced topic space, could the position of their *next* recalled event be predicted reliably? For each location in the reduced topic space, we computed the set of line segments connecting successively recalled events (across all participants) that intersected that location (see *Methods*). We then computed (for each location) the distribution of angles formed by the lines defined by those line segments and a fixed reference line (the *x*-axis). Rayleigh tests revealed the set of locations in topic space at which these across-participant distributions exhibited reliable peaks (blue arrows in Fig. 6B reflect significant peaks at *p* < 0.05, corrected). We observed that the locations traversed by nearly the entire episode trajectory exhibited such peaks. In other words, participants’ recalls exhibited similar trajectories to each other that also matched the trajectory of the original episode (Fig. 6C). This is especially notable when considering the fact that the number of HMM-identified recall events (dots in Fig. 6C) varied considerably across people, and that every participant used different words to describe what they had remembered happening in the episode. Differences in the numbers of recall events appear in participants’ trajectories as differences in the sampling resolution along the trajectory. We note that this framework also provides a means of disentangling classic “proportion recalled” measures (i.e., the proportion of episode events described in participants’ recalls) from participants’ abilities to recapitulate the episode’s essence (i.e., the similarity between the shapes of the original episode trajectory and that defined by each participant’s recounting of the episode).

In addition to enabling us to visualize the episode’s high-level essence, describing the episode as a geometric trajectory also enables us to drill down to individual words and quantify how each word relates to the memorability of each event. This provides another approach to examining participants’ recall for low-level details beyond the precision and distinctiveness measures we defined above. The results displayed in Figures 3C and 5A suggest that certain events were remembered better than others. Given this, we next asked asked whether the events that were generally remembered precisely or imprecisely tended to reflect particular content. Because our analysis framework projects the dynamic episode content and participants’ recalls into a shared space, and because the dimensions of that space represent topics (which are, in turn, sets of weights over known words in the vocabulary), we are able to recover the weighted combination of words that make up any point (i.e., topic vector) in this space. We first computed the average precision with which participants recalled each of the 30 episode events (Fig. 7A; note that this result is analogous to a serial position curve created from our precision metric). We then computed a weighted average of the topic vectors for each episode event, where the weights reflected how precisely each event was recalled. To visualize the result, we created a “wordle” image (Mueller et al., 2018) where words weighted more heavily by more precisely-remembered topics appear in a larger font (Fig. 7B, green box). Across the full episode, content that weighted heavily on topics and words central to the major foci of the episode (e.g., the names of the two main characters, “Sherlock” and “John,” and the address of a major recurring location, “221B Baker Street”) was best remembered. An analogous analysis revealed which themes were less-precisely remembered. Here in computing the weighted average over events’ topic vectors, we weighted each event in *inverse* proportion to its average precision (Fig. 7B, red box). The least precisely remembered episode content reflected information that was extraneous to the episode’s essence, such as the proper names of relatively minor characters (e.g., “Mike,” “Molly,” and “Lestrade”) and locations (e.g., “St. Bartholomew’s Hospital”).

**Figure 7:**
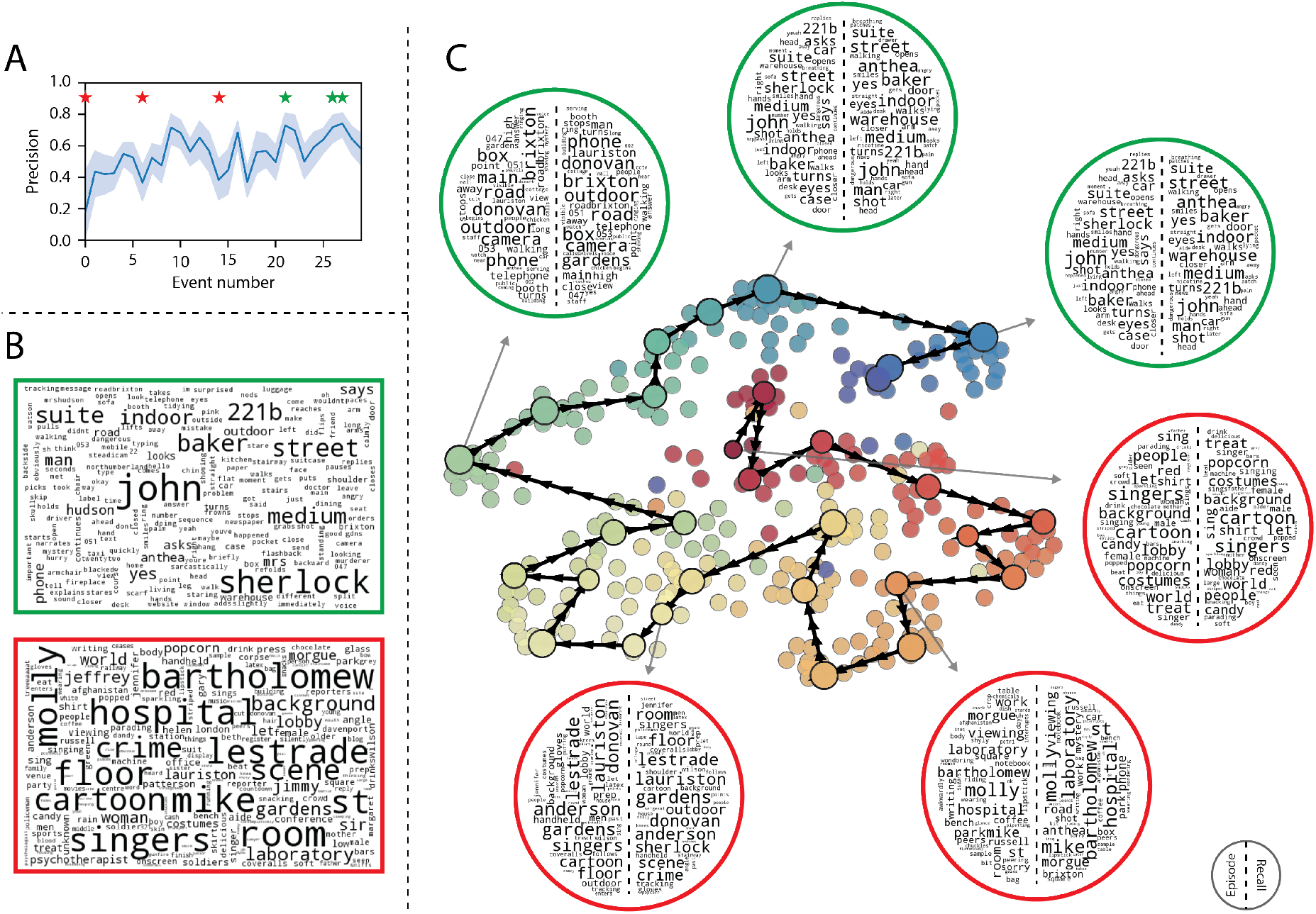
Language used in the most and least precisely remembered events. **A**. Average precision (episode event-recall event topic vector correlation) across participants for each episode event. Here we defined each episode event’s precision for each participant as the correlation between its topic vector and the most-correlated recall event’s topic vector from that participant. Error bars denote bootstrap-derived across-participant 95% confidence intervals. The stars denote the three most precisely remembered events (green) and least precisely remembered events (red). **B**. Wordles comprising the top 200 highest-weighted words reflected in the weighted-average topic vector across episode events. Green: episode events were weighted by their precision (Panel A). Red: episode events were weighted by the inverse of their precision. **C**. The set of all episode and recall events is projected onto the two-dimensional space derived in Figure 6. The dots outlined in black denote episode events (dot size is proportional to each event’s average precision). The dots without black outlines denote individual recall events from each participant. All dots are colored using the same scheme as Figure 6A. Wordles for several example events are displayed (green: three most precisely remembered events; red: three least precisely remembered events). Within each circular wordle, the left side displays words associated with the topic vector for the episode event, and the right side displays words associated with the (average) recall event topic vector, across all recall events matched to the given episode event.

A similar result emerged from assessing the topic vectors for individual episode and recall events (Fig. 7C). Here, for each of the three most and least precisely remembered episode events, we have constructed two wordles: one from the original episode event’s topic vector (left) and a second from the average recall topic vector for that event (right). The three most precisely remembered events (circled in green) correspond to scenes integral to the central plot-line: a mysterious figure spying on John in a phone booth; John meeting Sherlock at Baker St. to discuss the murders; and Sherlock laying a trap to catch the killer. Meanwhile, the least precisely remembered events (circled in red) reflect scenes that comprise minor plot points: a video of singing cartoon characters that participants viewed in an introductory clip prior to the main episode; John asking Molly about Sherlock’s habit of over-analyzing people; and Sherlock noticing evidence of Anderson’s and Donovan’s affair.

The results this far inform us about which aspects of the dynamic content in the episode participants watched were preserved or altered in participants’ memories. We next carried out a series of analyses aimed at understanding which brain structures might facilitate these preservations and transformations between the participants’ shared experience of watching the episode and their subsequent memories of the episode. In the first analysis, we sought to identify brain structures that were sensitive to the dynamic unfolding of the episode’s content, as characterized by its topic trajectory. We used a searchlight procedure to identify clusters of voxels whose activity patterns displayed a proximal temporal correlation structure (as participants watched the episode) matching that of the original episode’s topic proportions (Fig. 8A; see *Methods* for additional details). In a second analysis, we sought to identify brain structures whose responses (during episode viewing) reflected how each participant would later structure their *recounting* of the episode. We used a searchlight procedure to identify clusters of voxels whose proximal temporal correlation matrices matched that of the topic proportions matrix for each participant’s recall transcript (Figs. 8B; see *Methods* for additional details). To ensure our searchlight procedure identified regions *specifically* sensitive to the temporal structure of the episode or recalls (i.e., rather than those with a temporal autocorrelation length similar to that of the episode and recalls), we performed a phase shift-based permutation correction (see *Methods*). As shown in Figure 8C, the episode-driven searchlight analysis revealed a distributed network of regions that may play a role in processing information relevant to the narrative structure of the episode. The recall-driven searchlight analysis revealed a second network of regions (Fig. 8D) that may facilitate a person-specific transformation of one’s experience into memory. In identifying regions whose responses to ongoing experiences reflect how those experiences will be remembered later, this latter analysis extends classic *subsequent memory effect analyses* (e.g., Paller and Wagner, 2002) to the domain of naturalistic experiences.

**Figure 8:**
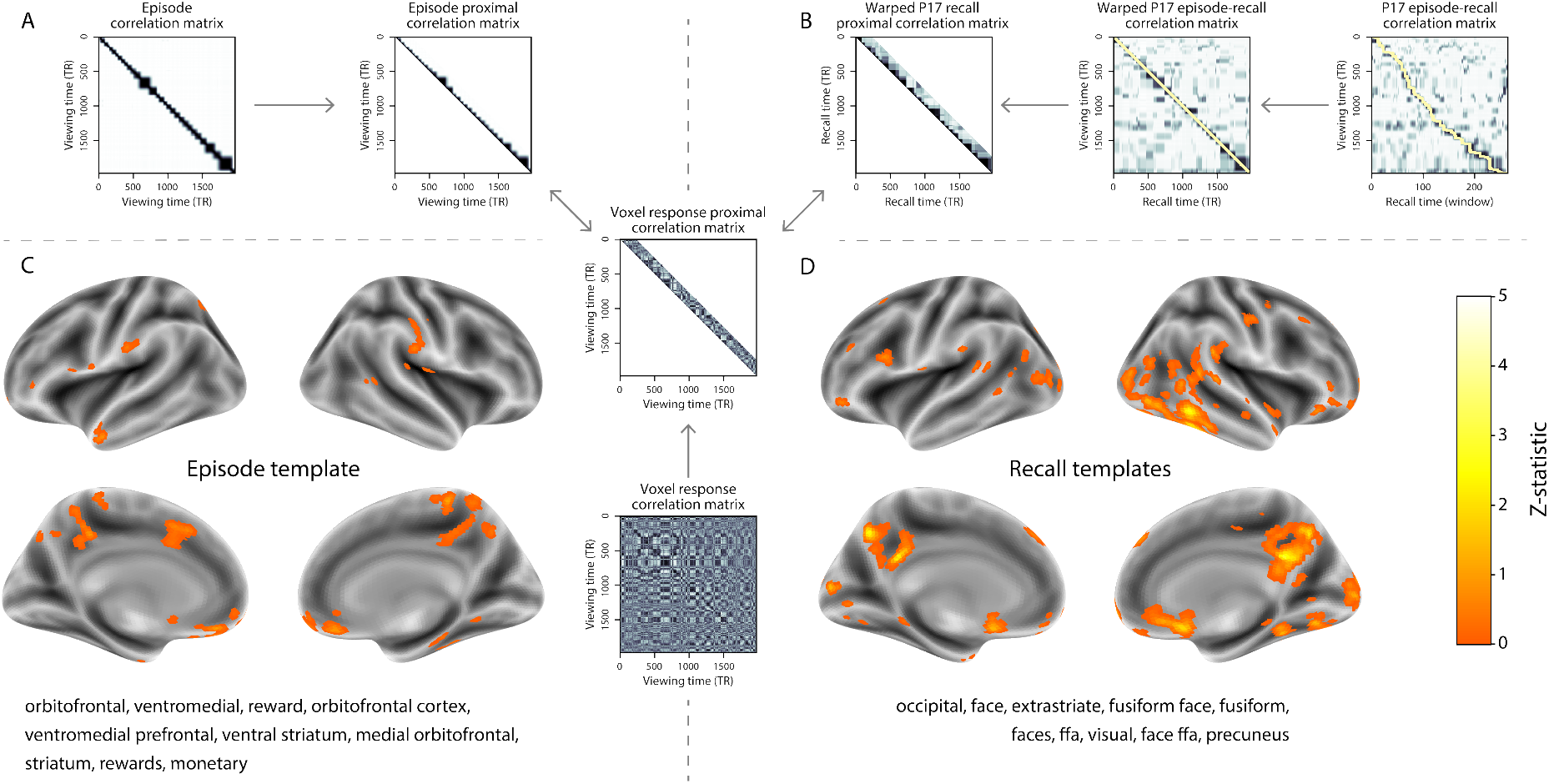
Brain structures that underlie the transformation of experience into memory. **A**. We isolated the proximal diagonals from the upper triangle of the episode correlation matrix, and applied this same diagonal mask to the voxel response correlation matrix for each cube of voxels in the brain. We then searched for brain regions whose activation timeseries consistently exhibited a similar proximal correlational structure to the episode model, across participants. **B**. We used dynamic time warping (Berndt and Clifford, 1994) to align each participant’s recall timeseries to the TR timeseries of the episode. We then applied the same diagonal mask used in Panel A to isolate the proximal temporal correlations and searched for brain regions whose activation timeseries for an individual consistently exhibited a similar proximal correlational structure to each individual’s recall. **C**. We identified a network of regions sensitive to the narrative structure of participants’ ongoing experience. The map shown is thresholded at *p* < 0.05, corrected. The top ten Neurosynth terms displayed in the panel were computed using the unthresholded map. **D**. We also identified a network of regions sensitive to how individuals would later structure the episode’s content in their recalls. The map shown is thresholded at *p* < 0.05, corrected. The top ten Neurosynth terms displayed in the panel were computed using the unthresholded map.

The searchlight analyses described above yielded two distributed networks of brain regions whose activity timecourses tracked with the temporal structure of the episode (Fig. 8C) or participants’ subsequent recalls (Fig. 8D). We next sought to gain greater insight into the structures and functional networks our results reflected. To accomplish this, we performed an additional, exploratory analysis using `Neurosynth` (Yarkoni et al., 2011). Given an arbitrary statistical map as input, `Neurosynth` performs a massive automated meta-analysis, returning a ranked list of terms frequently used in neuroimaging papers that report similar statistical maps. We ran `Neurosynth` on the (unthresholded) permutation-corrected maps for the episode- and recall-driven searchlight analyses. The top ten terms with maximally similar meta-analysis images identified by `Neurosynth` are shown in Figure 8.

## Discussion

Explicitly modeling the dynamic content of a naturalistic stimulus and participants’ memories enabled us to connect the present study of naturalistic recall with an extensive prior literature that has used list-learning paradigms to study memory (for review see Kahana, 2012), as in Figure 3. We found some similarities between how participants in the present study recounted a television episode and how participants typically recall memorized random word lists. However, our broader claim is that word lists miss out on fundamental aspects of naturalistic memory more like the sort of memory we rely on in everyday life. For example, there are no random word list analogs of character interactions, conceptual dependencies between temporally distant episode events, the sense of solving a mystery that pervades the *Sherlock* episode, or the myriad other features of the episode that convey deep meaning and capture interest. Nevertheless, each of these properties affects how people process and engage with the episode as they are watching it, and how they remember it later. The overarching goal of the present study is to characterize how the rich dynamics of the episode affect the rich behavioral and neural dynamics of how people remember it.

Our work casts remembering as reproducing (behaviorally and neurally) the topic trajectory, or “shape,” of an experience. When we characterized memory for a television episode using this framework, we found that every participant’s recounting of the episode recapitulated the low spatial frequency details of the shape of its trajectory through topic space (Fig. 6). We termed this narrative scaffolding the episode’s *essence*. Where participants’ behaviors varied most was in their tendencies to recount specific low-level details from each episode event. Geometrically, this appears as high spatial frequency distortions in participants’ recall trajectories relative to the trajectory of the original episode (Fig. 7). We developed metrics to characterize the precision (recovery of any and all event-level information) and distinctiveness (recovery of event-specific information). We also used word cloud visualizations to interpret the details of these event-level distortions.

The neural analyses we carried out (Fig. 8) also leveraged our geometric framework for characterizing the shapes of the episode and participants’ recountings. We identified one network of regions whose responses tracked with temporal correlations in the conceptual content of the episode (as quantified by topic models applied to a set of annotations about the episode). This network included orbitofrontal cortex, ventromedial prefrontal cortex, and striatum, among others. As reviewed by Ranganath and Ritchey (2012), several of these regions are members of the *anterior temporal system*, which has been implicated in assessing and processing the familiarity of ongoing experiences, emotions, social cognition, and reward. A second network we identified tracked with temporal correlations in the idiosyncratic conceptual content of participants’ subsequent recountings of the episode. This network included occipital cortex, extrastriate cortex, fusiform gyrus, and the precuneus. Several of these regions are members of the *posterior medial system* (Ranganath and Ritchey, 2012), which has been implicated in matching incoming cues about the current situation to internally maintained *situation models* that specify the parameters and expectations inherent to the current situation (also see Zacks et al., 2007; Zwaan and Radvansky, 1998). Taken together, our results support the notion that these two (partially overlapping) networks work in coordination to make sense of our ongoing experiences, distort them in a way that links them with our prior knowledge and experiences, and encodes those distorted representations into memory for our later use.

Our general approach draws inspiration from prior work aimed at elucidating the neural and behavioral underpinnings of how we process dynamic naturalistic experiences and remember them later. Our approach to identifying neural responses to naturalistic stimuli (including experiences) entails building an explicit model of the stimulus dynamics and searching for brain regions whose responses are consistent with the model (also see Huth et al., 2016, 2012). In prior work, a series of studies from Uri Hasson’s group (Baldassano et al., 2017; Chen et al., 2017; Lerner et al., 2011; Simony et al., 2016; Zadbood et al., 2017) have presented a clever alternative approach: rather than building an explicit stimulus model, these studies instead search for brain responses to the stimulus that are reliably similar across individuals. So called *inter-subject correlation* (ISC) and *inter-subject functional connectivity* (ISFC) analyses effectively treat other people’s brain responses to the stimulus as a “model” of how its features change over time (also see Simony and Chang, 2020). These purely brain-driven approaches are well suited to identifying which brain structures exhibit similar stimulus-driven responses across individuals. Further, because neural response dynamics are observed data (rather than model approximations), such approaches do not require a detailed understanding of which stimulus properties or features might be driving the observed responses. However, this also means that the specific stimulus features driving those responses are typically opaque to the researcher. Our approach is complementary. By explicitly modeling the stimulus dynamics, we are able to relate specific stimulus features to behavioral and neural dynamics. However, when our model fails to accurately capture the stimulus dynamics that are truly driving behavioral and neural responses, our approach necessarily yields an incomplete characterization of the neural basis of the processes we are studying.

Other recent work has used HMMs to discover latent event structure in neural responses to naturalistic stimuli (Baldassano et al., 2017). By applying HMMs to our explicit models of stimulus and memory dynamics, we gain a more direct understanding of those state dynamics. For example, we found that although the events comprising each participant’s recalls recapitulated the episode’s essence, participants differed in the *resolution* of their recounting of low-level details. In turn, these individual behavioral differences were reflected in differences in neural activity dynamics as participants watched the television episode.

Our approach also draws inspiration from the growing field of word embedding models. The topic models (Blei et al., 2003) we used to embed text from the episode annotations and participants’ recall transcripts are just one of many models that have been studied in an extensive literature. The earliest approaches to word embedding, including latent semantic analysis (Landauer and Dumais, 1997), used word co-occurrence statistics (i.e., how often pairs of words occur in the same documents contained in the corpus) to derive a unique feature vector for each word. The feature vectors are constructed so that words that co-occur more frequently have feature vectors that are closer (in Euclidean distance). Topic models are essentially an extension of those early models, in that they attempt to explicitly model the underlying causes of word co-occurrences by automatically identifying the set of themes or topics reflected across the documents in the corpus. More recent work on these types of semantic models, including word2vec (Mikolov et al., 2013), the Universal Sentence Encoder (Cer et al., 2018), GPT-2 (Radford et al., 2019), and GTP-3 (Brown et al., 2020) use deep neural networks to attempt to identify the deeper conceptual representations underlying each word. Despite the growing popularity of these sophisticated deep learning-based embedding models, we chose to prioritize interpretability of the embedding dimensions (e.g., Fig. 7) over raw performance (e.g., with respect to some predefined benchmark). Nevertheless, we note that our general framework is, in principle, robust to the specific choice of language model as well as other aspects of our computational pipeline. For example, the word embedding model, timeseries segmentation model, and the episode-recall matching function could each be customized to suit a particular question space or application. Indeed, for some questions, interpretability of the embeddings may not be a priority, and thus other text embedding approaches (including the deep learning-based models described above) may be preferable. Further work will be needed to explore the influence of particular models on our framework’s predictions and performance.

Our work has broad implications for how we characterize and assess memory in real-world settings, such as the classroom or physician’s office. For example, the most commonly used classroom evaluation tools involve simply computing the proportion of correctly answered exam questions. Our work indicates that this approach is only loosely related to what educators might really want to measure: how well did the students understand the key ideas presented in the course? Under this typical framework of assessment, the same exam score of 50% could be ascribed to two very different students: one who attended to the full course but struggled to learn more than a broad overview of the material, and one who attended to only half of the course but understood the attended material perfectly. Instead, one could apply our computational framework to build explicit dynamic content models of the course material and exam questions. This approach would provide a more nuanced and specific view into which aspects of the material students had learned well (or poorly). In clinical settings, memory measures that incorporate such explicit content models might also provide more direct evaluations of patients’ memories, and of doctor-patient interactions.

## Methods

### Paradigm and data collection

Data were collected by Chen et al. (2017). In brief, participants (*n* = 22) viewed the first 48 minutes of “A Study in Pink,” the first episode of the BBC television show *Sherlock*, while fMRI volumes were collected (TR = 1500 ms). Participants were pre-screened to ensure they had never seen any episode of the show before. The stimulus was divided into a 23 min (946 TR) and a 25 min (1030 TR) segment to mitigate technical issues related to the scanner. After finishing the clip, participants were instructed to (quoting from Chen et al., 2017) “describe what they recalled of the [episode] in as much detail as they could, to try to recount events in the original order they were viewed in, and to speak for at least 10 minutes if possible but that longer was better. They were told that completeness and detail were more important than temporal order, and that if at any point they realized they had missed something, to return to it. Participants were then allowed to speak for as long as they wished, and verbally indicated when they were finished (e.g., ‘I’m done’).” Five participants were dropped from the original dataset due to excessive head motion (2 participants), insufficient recall length (2 participants), or falling asleep during stimulus viewing (1 participant), resulting in a final sample size of *n* = 17. For additional details about the testing procedures and scanning parameters, see Chen et al. (2017). The testing protocol was approved by Princeton University’s Institutional Review Board.

After preprocessing the fMRI data and warping the images into a standard (3 mm^3^ MNI) space, the voxel activations were *z*-scored (within voxel) and spatially smoothed using a 6 mm (full width at half maximum) Gaussian kernel. The fMRI data were also cropped so that all episode-viewing data were aligned across participants. This included a constant 3 TR (4.5 s) shift to account for the lag in the hemodynamic response. (All of these preprocessing steps followed Chen et al., 2017, where additional details may be found.)

The video stimulus was divided into 1,000 fine-grained “time segments” and annotated by an independent coder. For each of these 1,000 annotations, the following information was recorded: a brief narrative description of what was happening, the location where the time segment took place, whether that location was indoors or outdoors, the names of all characters on-screen, the name(s) of the character(s) in focus in the shot, the name(s) of the character(s) currently speaking, the camera angle of the shot, a transcription of any text appearing on-screen, and whether or not there was music present in the background. Each time segment was also tagged with its onset and offset time, in both seconds and TRs.

### Data and code availability

The fMRI data we analyzed are available online here. The behavioral data and all of our analysis code may be downloaded here.

### Statistics

All statistical tests performed in the behavioral analyses were two-sided. All statistical tests performed in the neural data analyses were two-sided, except for the permutation-based thresholding, which was one-sided. In this case, we were specifically interested in identifying voxels whose activation time series reflected the temporal structure of the episode and recall topic proportions matrices to a *greater* extent than that of the phase-shifted matrices.

### Modeling the dynamic content of the episode and recall transcripts

#### Topic modeling

The input to the topic model we trained to characterize the dynamic content of the episode comprised 998 hand-generated annotations of short (mean: 2.96s) time segments spanning the video clip (Chen et al., 2017 generated 1000 annotations total; we removed two annotations referring to a break between the first and second scan sessions, during which no fMRI data were collected). We concatenated the text for all of the annotated features within each segment, creating a “bag of words” describing its content and performed some minor preprocessing (e.g., stemming possessive nouns and removing punctuation). We then re-organized the text descriptions into overlapping sliding windows spanning (up to) 50 annotations each. In other words, we estimated the “context” for each annotated segment using the text descriptions of the preceding 25 annotations, the present annotations, and the following 24 annotations. To model the context for annotations near the beginning of the episode (i.e., within 25 of the beginning or end), we created overlapping sliding windows that grew in size from one annotation to the full length. We also tapered the sliding window lengths at the end of the episode, whereby time segments within fewer than 24 annotations of the end of the episode were assigned sliding windows that extended to the end of the episode. This procedure ensured that each annotation’s content was represented in the text corpus an equal number of times.

We trained our model using these overlapping text samples with `scikit-learn` (version 0.19.1; Pedregosa et al., 2011), called from our high-dimensional visualization and text analysis software, `HyperTools` (Heusser et al., 2018b). Specifically, we used the `CountVectorizer` class to transform the text from each window into a vector of word counts (using the union of all words across all annotations as the “vocabulary,” excluding English stop words); this yielded a number-of-windows by number-of-words *word count* matrix. We then used the `LatentDirichletAllocation` class (topics=100, method=‘batch’) to fit a topic model (Blei et al., 2003) to the word count matrix, yielding a number-of-windows (1047) by number-of-topics (100) *topic proportions* matrix. The topic proportions matrix describes the gradually evolving mix of topics (latent themes) present in each annotated time segment of the episode. Next, we transformed the topic proportions matrix to match the 1976 fMRI volume acquisition times. We assigned each topic vector to the timepoint (in seconds) midway between the beginning of the first annotation and the end of the last annotation in its corresponding sliding text window. By doing so, we warped the linear temporal distance between consecutive topic vectors to align with the inconsistent temporal distance between consecutive annotations (whose durations varied greatly). We then rescaled these timepoints to 1.5s TR units, and used linear interpolation to estimate a topic vector for each TR. This resulted in a number-of-TRs (1976) by number-of-topics (100) matrix.

We created similar topic proportions matrices using hand-annotated transcripts of each participant’s verbal recall of the episode (annotated by Chen et al., 2017). We tokenized the transcript into a list of sentences, and then re-organized the list into overlapping sliding windows spanning (up to) 10 sentences each, analogously to how we parsed the episode annotations. In turn, we transformed each window’s sentences into a word count vector (using the same vocabulary as for the episode model), then used the topic model already trained on the episode scenes to compute the most probable topic proportions for each sliding window. This yielded a number-of-windows (range: 83–312) by number-of-topics (100) topic proportions matrix for each participant. These reflected the dynamic content of each participant’s recalls. Note: for details on how we selected the episode and recall window lengths and number of topics, see *Supporting Information* and Figure S1.

### Segmenting topic proportions matrices into discrete events using hidden Markov Models

We parsed the topic proportions matrices of the episode and participants’ recalls into discrete events using hidden Markov Models (HMMs; Rabiner, 1989). Given the topic proportions matrix (describing the mix of topics at each timepoint) and a number of states, *K*, an HMM recovers the set of state transitions that segments the timeseries into *K* discrete states. Following Baldassano et al. (2017), we imposed an additional set of constraints on the discovered state transitions that ensured that each state was encountered exactly once (i.e., never repeated). We used the `BrainIAK` toolbox (Capota et al., 2017) to implement this segmentation.

We used an optimization procedure to select the appropriate *K* for each topic proportions matrix. Prior studies on narrative structure and processing have shown that we both perceive and internally represent the world around us at multiple, hierarchical timescales (e.g., Baldassano et al., 2017, 2018; Chen et al., 2017; Hasson et al., 2015, 2008; Lerner et al., 2011). However, for the purposes of our framework, we sought to identify the single timeseries of event-representations that is emphasized *most heavily* in the temporal structure of the episode and of each participant’s recall. We quantified this as the set of *K* states that maximized the similarity between topic vectors for timepoints comprising each state, while minimizing the similarity between topic vectors for timepoints across different states. Specifically, we computed (for each matrix)

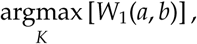

where *a* was the distribution of within-state topic vector correlations, and *b* was the distribution of across-state topic vector correlations. We computed the first Wasserstein distance (*W*_1_; also known as *Earth mover’s distance*; Dobrushin, 1970; Ramdas et al., 2017) between these distributions for a large range of possible *K*-values (range [2, 50]), and selected the *K* that yielded the maximum value. Figure 2B displays the event boundaries returned for the episode, and Figure S4 displays the event boundaries returned for each participant’s recalls. See Figure S6 for the optimization functions for the episode and recalls. After obtaining these event boundaries, we created stable estimates of the content represented in each event by averaging the topic vectors across timepoints between each pair of event boundaries. This yielded a number-of-events by number-of-topics matrix for the episode and recalls from each participant.

#### Naturalistic extensions of classic list-learning analyses

In traditional list-learning experiments, participants view a list of items (e.g., words) and then recall the items later. Our episode-recall event matching approach affords us the ability to analyze memory in a similar way. The episode and recall events can be treated analogously to studied and recalled “items” in a list-learning study. We can then extend classic analyses of memory performance and dynamics (originally designed for list-learning experiments) to the more naturalistic episode recall task used in this study.

Perhaps the simplest and most widely used measure of memory performance is *accuracy*—i.e., the proportion of studied (experienced) items (in this case, episode events) that the participant later remembered. Chen et al. (2017) used this method to rate each participant’s memory quality by computing the proportion of (50, manually identified) scenes mentioned in their recall. We found a strong across-participants correlation between these independent ratings and the proportion of 30 HMM-identified episode events matched to participants’ recalls (Pearson’s *r*(15) = 0.71, *p* = 0.002). We further considered a number of more nuanced memory performance measures that are typically associated with list-learning studies. We also provide a software package, `Quail`, for carrying out these analyses (Heusser et al., 2017).

#### Probability of first recall (PFR)

PFR curves (Atkinson and Shiffrin, 1968; Postman and Phillips, 1965; Welch and Burnett, 1924) reflect the probability that an item will be recalled first, as a function of its serial position during encoding. To carry out this analysis, we initialized a number-of-participants (17) by number-of-episode-events (30) matrix of zeros. Then, for each participant, we found the index of the episode event that was recalled first (i.e., the episode event whose topic vector was most strongly correlated with that of the first recall event) and filled in that index in the matrix with a 1. Finally, we averaged over the rows of the matrix, resulting in a 1 by 30 array representing the proportion of participants that recalled an event first, as a function of the order of the event’s appearance in the episode (Fig. 3A).

#### Lag conditional probability curve (lag-CRP)

The lag-CRP curve (Kahana, 1996) reflects the probability of recalling a given item after the just-recalled item, as a function of their relative encoding positions (or *lag*). In other words, a lag of 1 indicates that a recalled item was presented immediately after the previously recalled item, and a lag of -3 indicates that a recalled item came 3 items before the previously recalled item. For each recall transition (following the first recall), we computed the lag between the current recall event and the next recall event, normalizing by the total number of possible transitions. This yielded a number-of-participants (17) by number-of-lags (−29 to +29; 58 lags total excluding lags of 0) matrix. We averaged over the rows of this matrix to obtain a group-averaged lag-CRP curve (Fig. 3B).

#### Serial position curve (SPC)

SPCs (Murdock, 1962) reflect the proportion of participants that remember each item as a function of the item’s serial position during encoding. We initialized a number-of-participants (17) by number-of-episode-events (30) matrix of zeros. Then, for each recalled event, for each participant, we found the index of the episode event that the recalled event most closely matched (via the correlation between the events’ topic vectors) and entered a 1 into that position in the matrix. This resulted in a matrix whose entries indicated whether or not each event was recalled by each participant (depending on whether the corresponding entires were set to one or zero). Finally, we averaged over the rows of the matrix to yield a 1 by 30 array representing the proportion of participants that recalled each event as a function of the events’ order appearance in the episode (Fig. 3C).

#### Temporal clustering scores

Temporal clustering describes a participant’s tendency to organize their recall sequences by the learned items’ encoding positions. For instance, if a participant recalled the episode events in the exact order they occurred (or in exact reverse order), this would yield a score of 1. If a participant recalled the events in random order, this would yield an expected score of 0.5. For each recall event transition (and separately for each participant), we sorted all not-yet-recalled events according to their absolute lag (i.e., distance away in the episode). We then computed the percentile rank of the next event the participant recalled. We averaged these percentile ranks across all of the participant’s recalls to obtain a single temporal clustering score for the participant.

#### Semantic clustering scores

Semantic clustering describes a participant’s tendency to recall semantically similar presented items together in their recall sequences. Here, we used the topic vectors for each event as a proxy for its semantic content. Thus, the similarity between the semantic content for two events can be computed by correlating their respective topic vectors. For each recall event transition, we sorted all not-yet-recalled events according to how correlated the topic vector *of the closest-matching episode event* was to the topic vector of the closest-matching episode event to the just-recalled event. We then computed the percentile rank of the observed next recall. We averaged these percentile ranks across all of the participant’s recalls to obtain a single semantic clustering score for the participant.

#### Averaging correlations

In all instances where we performed statistical tests involving precision or distinctiveness scores (Fig. 5), we used the Fisher *z*-transformation (Fisher, 1925) to stabilize the variance across the distribution of correlation values prior to performing the test. Similarly, when averaging precision or distinctiveness scores, we *z*-transformed the scores prior to computing the mean, and inverse *z*-transformed the result.

#### Visualizing the episode and recall topic trajectories

We used the UMAP algorithm (McInnes et al., 2018) to project the 100-dimensional topic space onto a two-dimensional space for visualization (Figs. 6, 7). To ensure that all of the trajectories were projected onto the *same* lower dimensional space, we computed the low-dimensional embedding on a “stacked” matrix created by vertically concatenating the events-by-topics topic proportions matrices for the episode, across-participants average recall and all 17 individual participants’ recalls. We then separated the rows of the result (a total-number-of-events by two matrix) back into individual matrices for the episode topic trajectory, across-participant average recall trajectory, and the trajectories for each individual participant’s recalls (Fig. 6). This general approach for discovering a shared low-dimensional embedding for a collections of high-dimensional observations follows Heusser et al. (2018b).

We optimized the manifold space for visualization based on two criteria: First, that the 2D embedding of the episode trajectory should reflect its original 100-dimensional structure as faithfully as possible. Second, that the path traversed by the embedded episode trajectory should intersect itself a minimal number of times. The first criteria helps bolster the validity of visual intuitions about relationships between sections of episode content, based on their locations in the embedding space. The second criteria was motivated by the observed low off-diagonal values in the episode trajectory’s temporal correlation matrix (suggesting that the same topic-space coordinates should not be revisited; see Fig. 2A). For further details on how we created this low-dimensional embedding space, see *Supporting Information*.

#### Estimating the consistency of flow through topic space across participants

In Figure 6B, we present an analysis aimed at characterizing locations in topic space that different participants move through in a consistent way (via their recall topic trajectories). The two-dimensional topic space used in our visualizations (Fig. 6) comprised a 60 × 60 (arbitrary units) square. We tiled this space with a 50 × 50 grid of evenly spaced vertices, and defined a circular area centered on each vertex whose radius was two times the distance between adjacent vertices (i.e., 2.4 units). For each vertex, we examined the set of line segments formed by connecting each pair successively recalled events, across all participants, that passed through this circle. We computed the distribution of angles formed by those segments and the *x*-axis, and used a Rayleigh test to determine whether the distribution of angles was reliably “peaked” (i.e., consistent across all transitions that passed through that local portion of topic space). To create Figure 6B, we drew an arrow originating from each grid vertex, pointing in the direction of the average angle formed by the line segments that passed within 2.4 units. We set the arrow lengths to be inversely proportional to the *p*-values of the Rayleigh tests at each vertex. Specifically, for each vertex we converted all of the angles of segments that passed within 2.4 units to unit vectors, and we set the arrow lengths at each vertex proportional to the length of the (circular) mean vector. We also indicated any significant results (*p* < 0.05, corrected using the Benjamani-Hochberg procedure) by coloring the arrows in blue (darker blue denotes a lower *p*-value, i.e., a longer mean vector); all tests with *p* ≥ 0.05 are displayed in gray and given a lower opacity value.

### Searchlight fMRI analyses

In Figure 8, we present two analyses aimed at identifying brain regions whose responses (as participants viewed the episode) exhibited a particular temporal structure. We developed a searchlight analysis wherein we constructed a 5 × 5 × 5 cube of voxels (following Chen et al., 2017) centered on each voxel in the brain, and for each of these cubes, computed the temporal correlation matrix of the voxel responses during episode viewing. Specifically, for each of the 1976 volumes collected during episode viewing, we correlated the activity patterns in the given cube with the activity patterns (in the same cube) collected during every other timepoint. This yielded a 1976 × 1976 correlation matrix for each cube. Note: participant 5’s scan ended 75s early, and in Chen et al. (2017)’s publicly released dataset, their scan data was zero-padded to match the length of the other participants’. For our searchlight analyses, we removed this padded data (i.e., the last 50 TRs), resulting in a 1925 × 1925 correlation matrix for each cube in participant 5’s brain.

Next, we constructed a series of “template” matrices. The first template reflected the time-course of the episode’s topic proportions matrix, and the others reflected the timecourse of each participant’s recall topic proportions matrix. To construct the episode template, we computed the correlations between the topic proportions estimated for every pair of TRs (prior to segmenting the topic proportions matrices into discrete events; i.e., the correlation matrix shown in Figs. 2B and 8A). We constructed similar temporal correlation matrices for each participant’s recall topic proportions matrix (Figs. 2D, S4). However, to correct for length differences and potential non-linear transformations between viewing time and recall time, we first used dynamic time warping (Berndt and Clifford, 1994) to temporally align participants’ recall topic proportions matrices with the episode topic proportions matrix. An example correlation matrix before and after warping is shown in Fig. 8B. This yielded a 1976 × 1976 correlation matrix for the episode template and for each participant’s recall template.

The temporal structure of the episode’s content (as described by our model) is captured in the block-diagonal structure of the episode’s temporal correlation matrix (e.g., Figs. 2B, 8A), with time periods of thematic stability represented as dark blocks of varying sizes. Inspecting the episode correlation matrix suggests that the episode’s semantic content is highly temporally specific (i.e., the correlations between topic vectors from distant timepoints are almost all near zero). By contrast, the activity patterns of individual (cubes of) voxels can encode relatively limited information on their own, and their activity frequently contributes to multiple separate functions (Charron and Koechlin, 2010; Freedman et al., 2001; Rishel et al., 2013; Sigman and Dehaene, 2008). By nature, these two attributes give rise to similarities in activity across large timescales that may not necessarily reflect a single task. To enable a more sensitive analysis of brain regions whose shifts in activity patterns mirrored shifts in the semantic content of the episode or recalls, we restricted the temporal correlations we considered to the timescale of semantic information captured by our model. Specifically, we isolated the upper triangle of the episode correlation matrix and created a “proximal correlation mask” that included only diagonals from the upper triangle of the episode correlation matrix up to the first diagonal that contained no positive correlations. Applying this mask to the full episode correlation matrix was equivalent to excluding diagonals beyond the corner of the largest diagonal block. In other words, the timescale of temporal correlations we considered corresponded to the longest period of thematic stability in the episode, and by extension the longest period of thematic stability in participants’ recalls and the longest period of stability we might expect to see in voxel activity arising from processing or encoding episode content. Figure 8 shows this proximal correlation mask applied to the temporal correlation matrices for the episode, an example participant’s (warped) recall, and an example cube of voxels from our searchlight analyses.

To determine which (cubes of) voxel responses matched the episode template, we correlated the proximal diagonals from the upper triangle of the voxel correlation matrix for each cube with the proximal diagonals from episode template matrix (Kriegeskorte et al., 2008). This yielded, for each participant, a voxelwise map of correlation values. We then performed a one-sample *t*-test on the distribution of (Fisher *z*-transformed) correlations at each voxel, across participants. This resulted in a value for each voxel (cube), describing how reliably its timecourse followed that of the episode.

We further sought to ensure that our analysis identified regions where the activations’ temporal structure specifically reflected that of the episode, rather than regions whose activity was simply autocorrelated at a timescale similar to the episode template’s diagonal. To achieve this, we used a phase shift-based permutation procedure, whereby we circularly shifted the episode’s topic proportions matrix by a random number of timepoints (rows), computed the resulting “null” episode template, and re-ran the searchlight analysis, in full. (For each of the 100 permutations, the same random shift was used for all participants). We *z*-scored the observed (unshifted) result at each voxel against the distribution of permutation-derived “null” results, and estimated a *p*-value by computing the proportion of shifted results that yielded larger values. To create the map in Figure 8C, we thresholded out any voxels whose similarity to the unshifted episode’s structure fell below the 95^th^ percentile of the permutation-derived similarity results.

We used an analogous procedure to identify which voxels’ responses reflected the recall templates. For each participant, we correlated the proximal diagonals from the upper triangle of the correlation matrix for each cube of voxels with the proximal diagonals from the upper triangle of their (time-warped) recall correlation matrix. As in the episode template analysis, this yielded a voxelwise map of correlation coefficients for each participant. However, whereas the episode analysis compared every participant’s responses to the same template, here the recall templates were unique for each participant. As in the analysis described above, we *t*-scored the (Fisher *z*-transformed) voxelwise correlations, and used the same permutation procedure we developed for the episode responses to ensure specificity to the recall timeseries and assign significance values. To create the map in Figure 8D we again thresholded out any voxels whose scores were below the 95^th^ percentile of the permutation-derived null distribution.

#### Neurosynth decoding analyses

`Neurosynth` parses a massive online database of over 14,000 neuroimaging studies and constructs meta-analysis images for over 13,000 psychology- and neuroscience-related terms, based on NIfTI images accompanying studies where those terms appear at a high frequency. Given a novel image (tagged with its value type; e.g., *z*-, *t*-, *F*- or *p*-statistics), `Neurosynth` returns a list of terms whose meta-analysis images are most similar. Our permutation procedure yielded, for each of the two searchlight analyses, a voxelwise map of *z*-values. These maps describe the extent to which each voxel *specifically* reflected the temporal structure of the episode or individuals’ recalls (i.e., relative to the null distributions of phase-shifted values). We inputted the two statistical maps described above to `Neurosynth` to create a list of the 10 most representative terms for each map.

## Supporting information

Supporting information is available in the online version of the paper.

## Acknowledgements

We thank Luke Chang, Janice Chen, Chris Honey, Lucy Owen, Emily Whitaker, and Kirsten Ziman for feedback and scientific discussions. We also thank Janice Chen, Yuan Chang Leong, Kenneth Norman, and Uri Hasson for sharing the data used in our study. Our work was supported in part by NSF EPSCoR Award Number 1632738. The content is solely the responsibility of the authors and does not necessarily represent the official views of our supporting organizations.

## Author contributions

Conceptualization: A.C.H. and J.R.M.; Methodology: A.C.H., P.C.F. and J.R.M.; Software: A.C.H., P.C.F. and J.R.M.; Analysis: A.C.H., P.C.F. and J.R.M.; Writing, Reviewing, and Editing: A.C.H., P.C.F. and J.R.M.; Supervision: J.R.M.

## Author information

The authors declare no competing financial interests.

## Notes

https://github.com/ContextLab/sherlock-topic-model-paper

### Competing Interest Statement

The authors have declared no competing interest.

### Summary of Updates

Major refactor of introduction and discussion, methods overhaul

