## Supporting Information for "Geometric models reveal behavioral and neural signatures of transforming naturalistic experiences into episodic memories"

September 4, 2020

### Overview

This document provides additional details about the methods we used in the main text. We also include some additional analyses referenced in the main text.

### Additional details about topic modeling methods and results

#### Optimizing topic model parameters

In order to create accurate episode and recall models, we used an optimization method that was driven by our ability to explain hand-annotated memory performance metrics collected by Chen et al. (2017). In an earlier variant of our study (Heusser and Manning, 2018), we used a grid search to compute the $\omega$ (episode sliding window duration, in scenes), $\rho$ (recall sliding window duration, in sentences), and $K$ (number of topics) that satisfied

$$\underset{\omega,\rho,K}{\operatorname{argmax}} \left[\operatorname{corr}\left(\operatorname{corr}\left(\mu\left(\omega,\rho,K\right),\nu\left(\omega,\rho,K\right)\right),\theta\right)\right],$$

where corr($\mu,\nu$) is the per-participant correlation between the episode ($\mu$) and recall ($\nu$) topic proportions matrices, and $\theta$ is the per-participant hand-counted number of recalled scenes. We searched over a grid of pre-specified values for each of these parameters; the resulting correlations are displayed in Figure S1. The optimal parameters were $\omega = 50$, $\rho = 10$, and $K = 100$. In our current paper we made a number of improvements to how we preprocessed text and fit topic models (see *Methods*), but we carried the same optimal parameters forward from Heusser and Manning (2018) without performing any additional optimization.

The optimized model converged on 32 unique topics that were assigned non-zero weights. We provide a list of the top ten highest-weighted words from each topic in Figure S2.

#### Feature importance analyses

To determine the contribution of each feature to the temporal structure of the episode topic proportions matrix, we conducted a "leave one out" analysis. Specifically, we compared the original

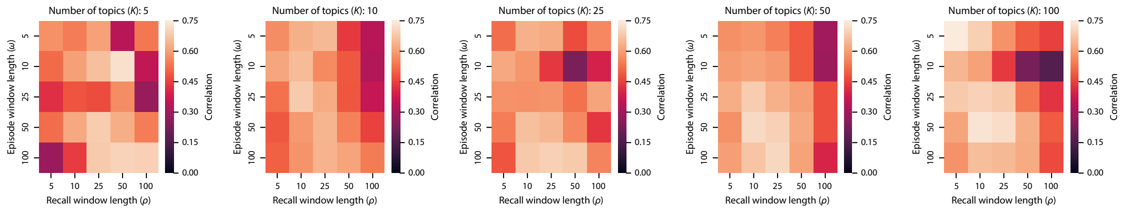


**Figure S1: Optimizing topic model parameters.** We performed a grid search over episode sliding window length ($\omega \in \{5, 10, 25, 50, 100\}$), recall sliding window length ($\rho \in \{5, 10, 25, 50, 100\}$), and number of topics ($K \in \{5, 10, 25, 50, 100\}$). The reported correlations are between per-participant episode-recall correlations and per-participant hand-counted numbers of recalled scenes.

| Topic ID | Top 10 words | Topic description |
| --- | --- | --- |
| 1 | john, outdoor, yes, phone, road, brixton, box, medium, donovan, street | John being watched |
| 2 | sherlock, john, indoor, laboratory, hospital, st, bartholomew, medium, yes, mike | Sherlock and John meet |
| 4 | man, john, warehouse, indoor, yes, medium, shoulder, says, hand, asks | Meeting with Mycroft |
| 5 | john, mike, sherlock, medium, molly, park, russell, square, outdoor, bench | Running into an old friend |
| 7 | yes, jeffrey, sir, jimmy, indoor, aide, medium, helen, woman, gary | First and second murder |
| 9 | sherlock, floor, room, crime, scene, lauriston, indoor, gardens, john, yes | Examining the body (a) |
| 17 | sherlock, lestrade, john, indoor, gardens, lauriston, room, medium, floor, scene | Lestrade calls in Sherlock (a) |
| 20 | soldiers, singers, cartoon, background, medium, indoor, world, yes, afghanistan, lobby | Intro cartoon/War flashback |
| 22 | sherlock, john, street, baker, 221b, indoor, mrs, hudson, suite, yes | 221B Baker St. (a) |
| 27 | sherlock, john, outdoor, medium, taxi, road, yes, says, phone, lauriston | Sherlock analyzes John (a) |
| 28 | sherlock, john, lestrade, lauriston, gardens, medium, anderson, donovan, indoor, yes | Who was Rachel? |
| 30 | sherlock, john, indoor, mediu, baker, street, 221b, suite, yes, phone | John texts the killer |
| 32 | donovan, lestrade, indoor, medium, aide, jimmy, press, room, conference, yes | Press conference (a) |
| 35 | john, sherlock, street, baker, medium, says, indoor, mrs, hudson, sequence | John leaves the flat |
| 37 | sherlock, john, street, suite, 221b, baker, indoor, medium, says, asks | 221B Baker St. (b) |
| 40 | sherlock, lestrade, yes, room, gardens, indoor, lauriston, floor, john, scene | Examining the body (b) |
| 42 | sherlock, molly, johm, bartholomew, st, hospital, medium, indoor, mike, room | John meets Sherlock Holmes |
| 46 | singers, cartoon, background, indoor, world, lobby, popcorn, yes, people, medium | Intro cartoon |
| 48 | john, donovan, gardens, lauriston, yes, street, outdoor, medium, shoulder, policeman | Unpopular with the police (a) |
| 65 | lestrade, donovan, indoor, room, press, conference, police, reporter, medium, reporters | Press conference (b) |
| 68 | john, medium, anthea, yes, indoor, street, baker, sherlock, outdoor, man | Kidnapping John |
| 70 | john, man, yes, warehouse, indoor, medium, shoulder, says, anthea, continues | Anthea brings John to Mycroft |
| 72 | sherlock, john, lestrade, lauriston, gardens, medium, indoor, yes, stairway, stairs | Lestrade calls in Sherlock (b) |
| 73 | jimmy, yes, sir, jeffrey, indoor, medium, gary, psychotherapist, helen, john | John's psychotherapist (a) |
| 75 | sherlock, john, donovan, outdoor, medium, gardens, lauriston, street, anderson, says | Unpopular with the police (b) |
| 78 | john, lestrade, mike, medium, donovan, indoor, park, room, conference, press | John takes a walk |
| 80 | john, sherlock, yes, indoor, laboratory, bartholomew, st, hospital, medium, mike | Sherlock analyzes John (b) |
| 84 | john, anthea, yes, road, man, outdoor, phone, medium, brixton, car | Anthea drives John home |
| 85 | sherlock, john, yes, taxi, outdoor, road, medium, says, phone, continues | Chasing the killer |
| 87 | john, indoor, jeffrey, sir, psychotherapist, medium, yes, room, london, helen | John's psychotherapist (b) |
| 91 | sherlock, john, lestrade, room, floor, crime, scene, gardens, lauriston, indoor | Examining the body (c) |
| 93 | john, room, indoor, medium, soldiers, psychotherapist, yes, outdoor, close, soldier | War flashback |

**Figure S2: Topics discovered in *Sherlock*.** We applied a topic model to hand-annotated information about 1000 scenes spanning the 48-minute episode. We identified 32 unique topics with non-zero weights (we used $K = 100$ topics to fit the model). Each topic comprises a distribution of weights over all words in the vocabulary. For each topic, we show the words with the 10 largest weights, along with a suggested description of the topic.

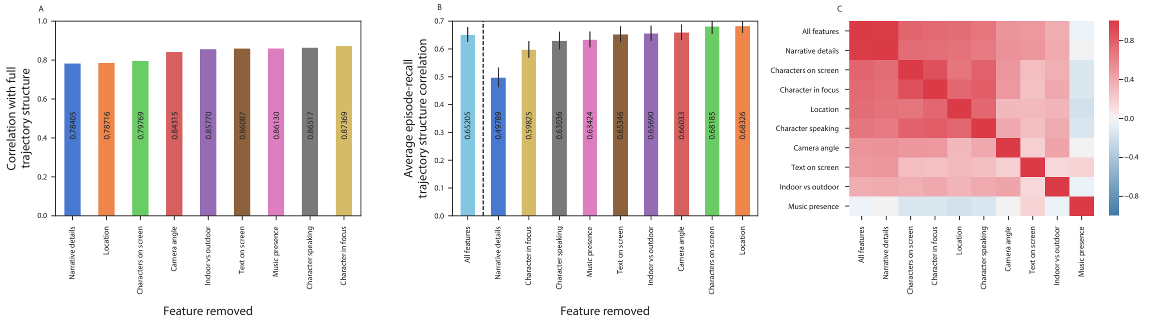


**Figure S3: Feature importance analysis.** **A.** Contributions of each feature type to the structure of the episode topic proportions matrix. The bar heights reflect the correlation between the proximal temporal structure (see *Methods*) of the episode topic proportions matrix computed using all features, and that of an episode topic proportions matrix computed using all features except the indicated feature. Lower bars reflect features that contribute more substantially to the episode's temporal structure. **B.** Which features are preserved during recall? The bar heights reflect the (average) across-participant correlations between the proximal temporal structure of episode and recall topic proportions matrices constructed in the absence of the given feature. Error bars denote standard error of the mean. **C.** Feature correlation matrix. Each entry displays the correlation between episode topic portions matrices created using only the indicated (row/column) features.

episode's topic proportions matrix (created using all hand-annotated features from the 998 manually identified scenes spanning the *Sherlock* episode; see *Methods* for a full list of features) with topic proportions matrices created using all but one type of feature. For each impoverished topic proportions matrix, we computed the timepoint-by-timepoint correlation matrix, and correlated the proximal diagonals from the upper triangle with those from the temporal correlation matrix of the feature-complete matrix (for details on how we isolated proximal temporal correlations, see *Methods*). Observing a lower correlation between an impoverished matrix (with a particular feature removed) and the feature-complete matrix would suggest that the held-out feature contributed more prominently to the full episode topic proportion matrix's temporal structure. We found that hand-annotated narrative details played the greatest roll in determining the temporal structure of the episode, whereas the name of the character(s) in focus for each shot contributed least (Fig. S3A).

Next, we sought to determine which annotated features contributed aspects of the episode's temporal structure that were preserved in participants' later recalls. Specifically, we computed the timepoint-by-timepoint correlation matrix of the episode's topic proportions matrix, and correlated the proximal diagonals from its upper triangle with those from the timepoint-by-timepoint correlation matrices for each participant's recall topic proportions matrices (stretched via linear interpolation to have the same number of timepoints as the episode topic proportions matrix). This yielded a single correlation coefficient for each participant. We then repeated this analysis with each annotated feature held out in turn. Observing a lower correlation between the episode and recall topic proportions matrices (constructed in the absence of a given feature) would indicate that participants utilize changes in that feature's content to discriminate between sections of the episode when organizing their recalls. We found that hand-annotated narrative details were the most heavily utilized feature, whereas changes in the text present on-screen, the indoor/outdoor distinction between shots, the camera angle, the names of the various characters on screen, and the shot's location tended not to impact participants' recall structures (i.e., removing those features resulted in a *greater* episode-recall correlation than including them; Fig. S3B).

We also wondered how the different types of features might relate. For example, knowing which character is in focus during a given scene may also provide information about which character is speaking. We computed topic proportions matrices from the annotations for each individual feature, in turn, and (using the same technique as in the above analyses) compared the proximal temporal correlation structure of each single-feature topic proportions matrix to each other, as well as to that of the full episode. This provided additional confirmation that the full episode's temporal structure was largely driven by narrative details. We also found that character-driven features (characters on screen, characters speaking, and characters in focus) were strongly correlated. Other details, such as the presence or absence of music, led to very different topic proportions matrices (Fig. S3C).

#### Creating a low-dimensional embedding space

Figures 6 and 7C in the main text display two-dimensional projections of the 100-dimensional topic trajectories for the episode (Figs. 6A, 7C), average recall (Fig. 6B), and each individual's recall (Figs. 6C, 7C). We created these embeddings using the Uniform Manifold Approximation and Projection algorithm (UMAP; McInnes et al., 2018) called from our high-dimensional visualization and text analysis software, `HyperTools` (Heusser et al., 2018). An advantage of the UMAP algorithm over comparable manifold learning techniques (e.g., *t*-SNE) is that UMAP explicitly attempts to preserve the global structure of the data (McInnes et al., 2018; Becht et al., 2019) by constructing a space where distance on the manifold is standard Euclidean distance, with respect to the global coordinate system. This was important in our use case, as we wanted to visualize both the evolving structure of the episode and the spatial relationships between presented and recalled content.

UMAP achieves a balance between representing local and global structure via a subset of its hyperparameters: `n_neighbors` , `spread`, and `min_dist`. The `n_neighbors` hyperparameter ($K$) denotes the number of nearest neighbors to consider in constructing the high-dimensional fuzzy simplicial set for each datapoint. The `spread` ($\gamma$) and `min_dist` ($\delta$) hyperparameters function together to create the differentiable decay curve used to approximate the injective function for mapping between high- and low-dimensional fuzzy simplicial sets. In brief, `min_dist` determines the degree to which nearby points are clustered or expanded, relative to to the overall `spread`.

Two other parameters ultimately affect this balance between preserving local versus global structure: the seed ($\tau$) for the (pseudo-)random number generator (RNG), and the order of the observations (i.e., trajectories) to be embedded ($\varphi$). As described in *Methods*, we ensured the episode and recall events were projected onto the *same* low-dimensional manifold by fitting the embedding model to a stacked matrix of all episode, average recall, and individuals' recall events. After initializing the low-dimensional simplicial set (by default, using a spectral embedding of the high-dimensional simplicial set's fuzzy 1-skeleton), UMAP optimizes the embedding using stochastic gradient descent with cross-entropy as a cost function. During optimization, indices of the data are sampled at random, and thus the local minimum achieved by the optimization is dependent on both the state of the RNG and the sequence in which observations are concatenated.

We performed a grid search over pre-specified values of these hyperparameters, and optimized the manifold space for visualization based on two criteria. First, the 2D embedding of the episode trajectory should reflect its original 100-dimensional structure as faithfully as possible. Second, the path traversed by the embedded episode trajectory should intersect itself a minimal number of times. The first criteria helps bolster the validity of visual intuitions about relationships between sections of episode content, based on their locations in the embedding space. The second criteria was motivated by the observed low off-diagonal values in the episode topic proportions matrix's temporal correlation matrix (suggesting that the same topic-space coordinates should not be re-visited; see Figure 2A in the main text). Further, we found through simulation that our statistical procedure for testing the consistency of trajectory directions across participants (Fig. 6B, also see *Methods*) can be confounded when and where the topic trajectory intersects itself. We formalized this optimization as the combination of hyperparameter values that satisfied

$$\underset{\{K,\gamma,\delta,\tau,\varphi \,\in\, E \,\mid\, \Phi(K,\gamma,\delta,\tau,\varphi)\}}{\operatorname{argmax}} \left[\operatorname{corr}\left(\Upsilon\left(\xi_{K,\gamma,\delta,\tau,\varphi}\right),\ \Upsilon\left(\chi\right)\right)\right], \text{ where}$$

$$\Phi\left(K,\gamma,\delta,\tau,\varphi\right) = \underset{K,\gamma,\delta,\tau,\varphi}{\operatorname{argmin}}\left[\Gamma\left(\xi_{K,\gamma,\delta,\tau,\varphi}\right)\right],$$

$\xi$ is the episode's trajectory through the manifold space; $\chi$ is the original, 100-dimensional episode trajectory; $\Upsilon$ is a function that computes a condensed matrix of pairwise distances between event vectors (computed using correlation distance in the original topic space and Euclidean distance in the manifold space); $\operatorname{corr}\left(\Upsilon\left(\xi\right),\Upsilon\left(\chi\right)\right)$ is the correlation between the sets of pairwise distances, and $\Gamma$ is the number of instances in which lines drawn between consecutive episode event embeddings intersected each other. The sets of hyperparameter values we searched over comprised: $K \in \{10x \in \mathbb{Z} \mid 10 < x < 22\} \cup \{161\}$ (a range roughly centered on half the total number of events, 161, in order to balance representations of local and global structure); $\gamma \in \{1,3,5,7,9\}$; $\delta \in \{0.1,0.3,0.5,0.7,0.9\}$; $\tau \in \{x \in \mathbb{Z} \mid 0 < x < 101\}$; and $\varphi \in \binom{S}{3}$, where $S = \{\text{episode events, average recall events, individual recall events}\}$. The optimal parameters (that yielded $\Phi = 0$) were $K = 170$, $\gamma = 7$, $\delta = 0.7$, $\tau = 41$, with the order of sequence $\varphi$ as the average recall events, episode events, and individuals' recall events, vertically concatenated, in order.

#### Participant-level figures referenced in the main text

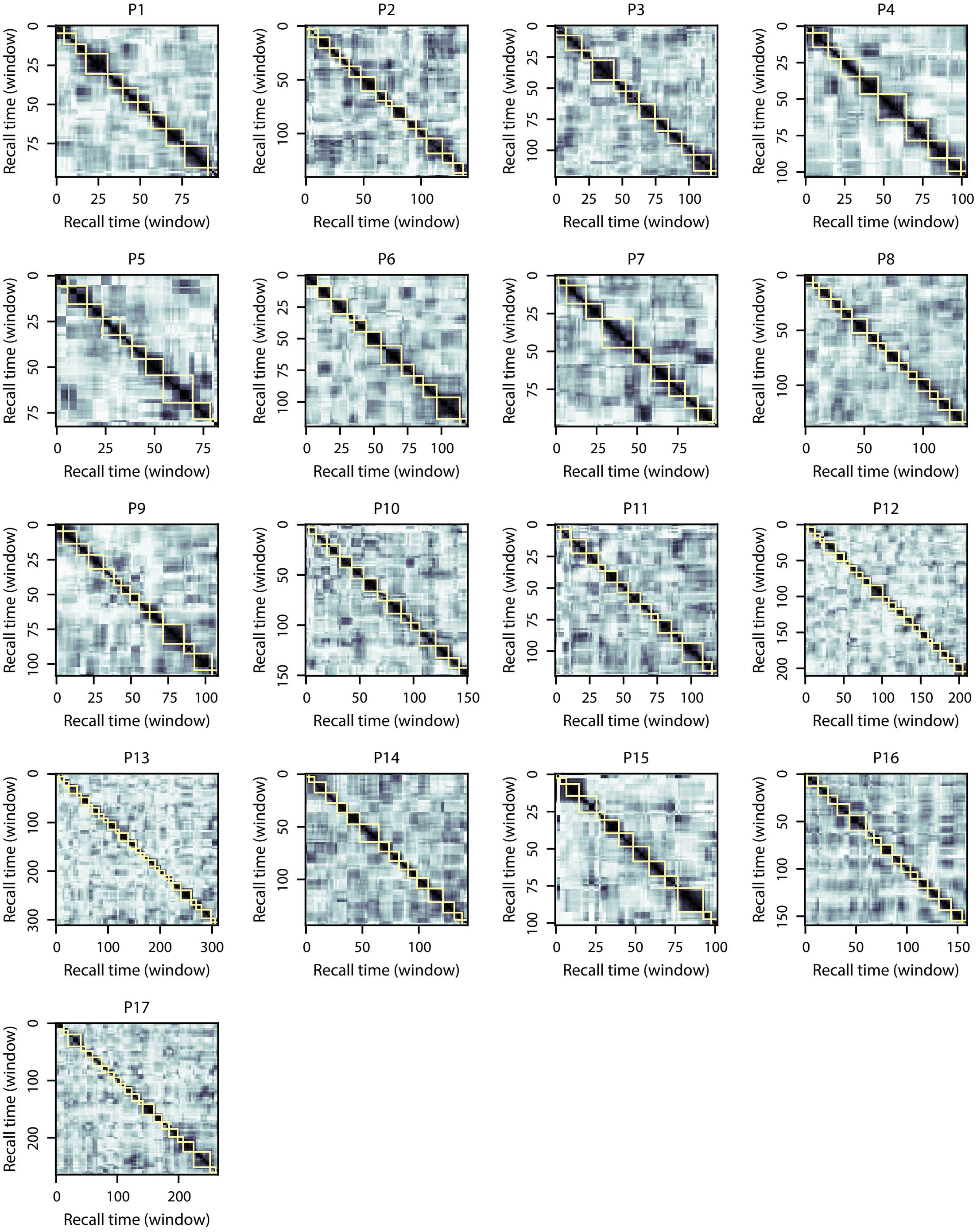


**Figure S4: Recall temporal correlation matrices and event segmentation fits.** Each panel is in the same format as Figure 2E in the main text. The yellow boxes indicate HMM-identified event boundaries.

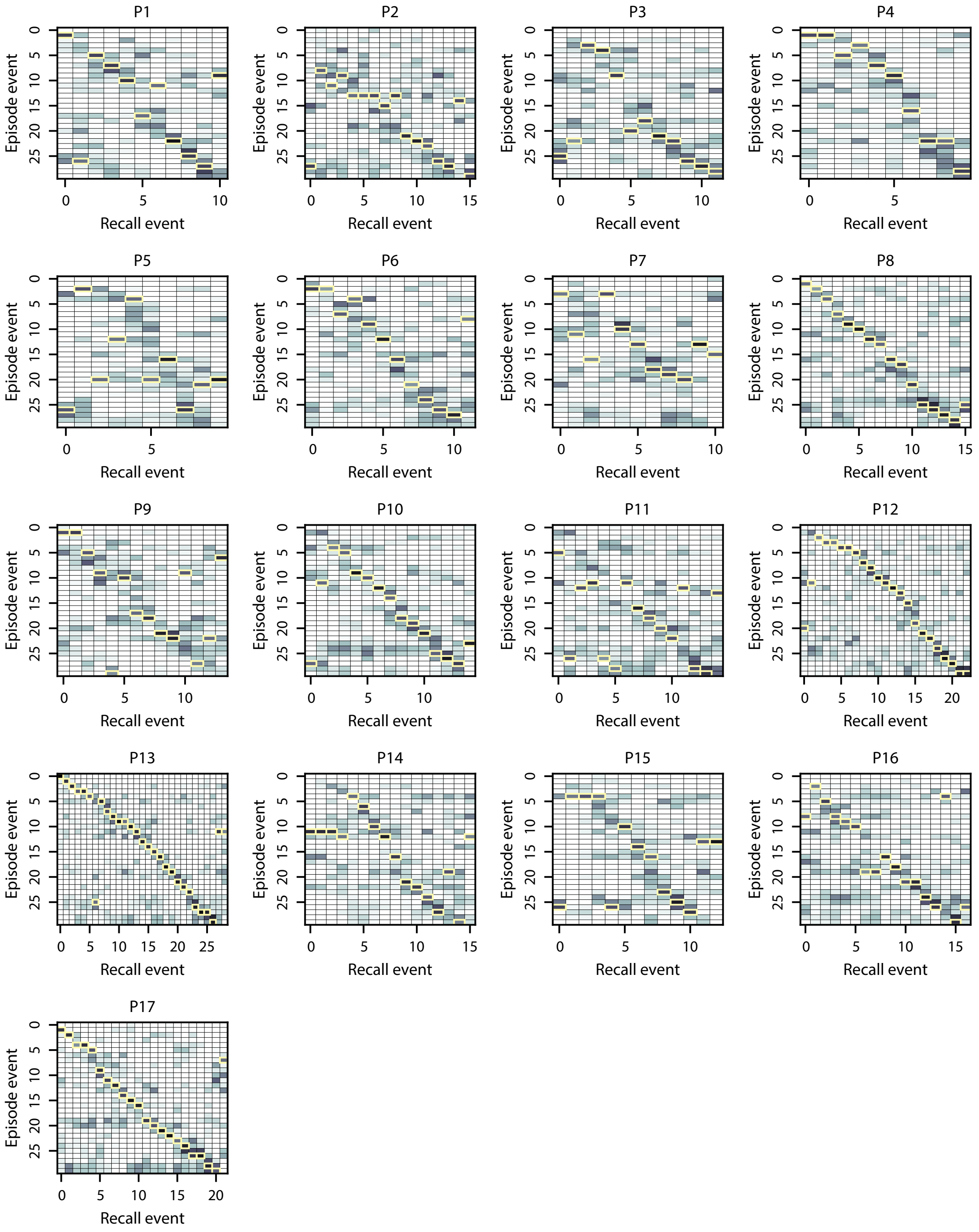


**Figure S5: Episode-recall event correlation matrices.** Each panel is in the same format as Figure 2G in the main text. The yellow boxes mark the matched episode event for each recall event (i.e., the maximum correlation in each column).

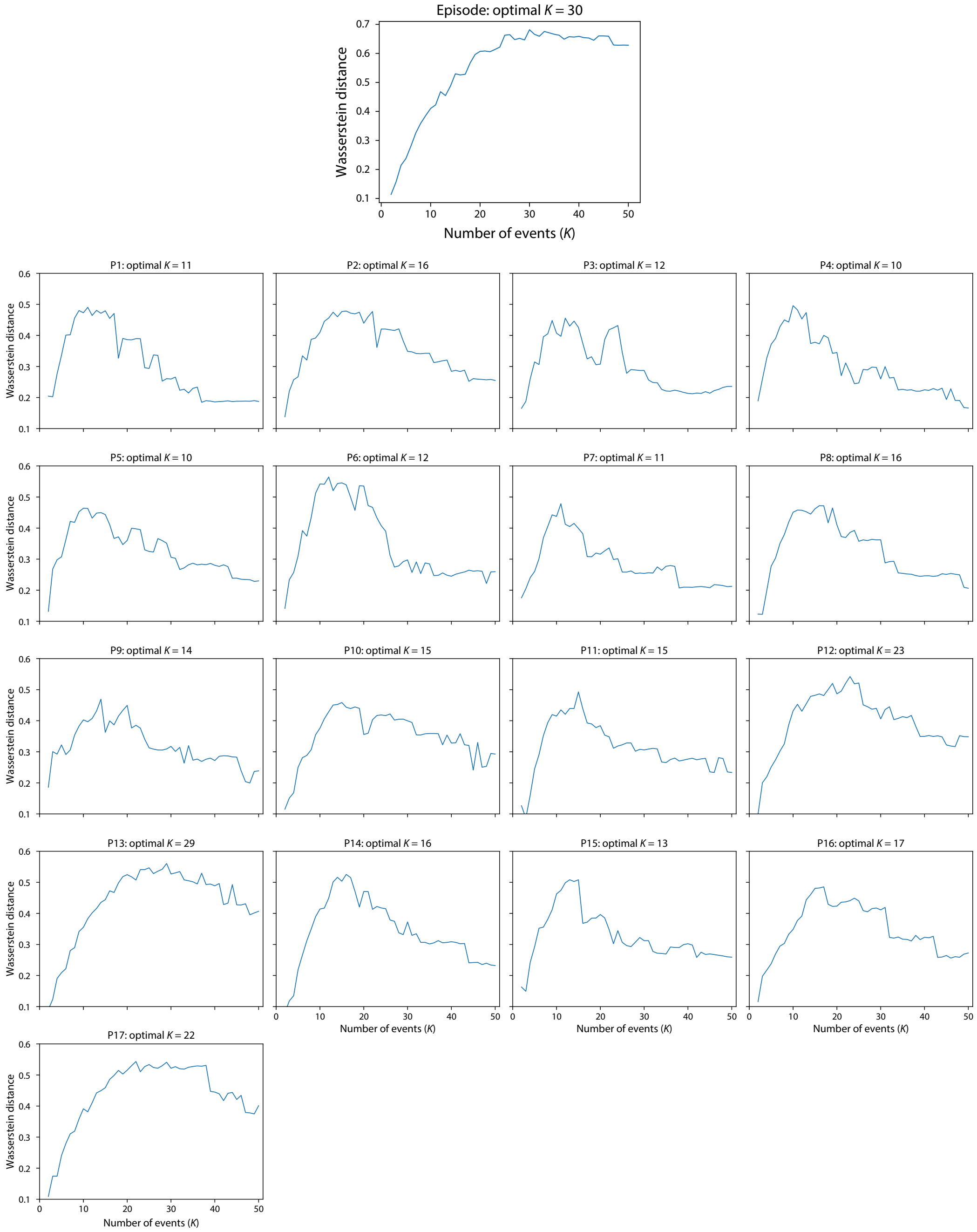


**Figure S6: Episode and recall topic proportions matrix $K$-optimization functions.** We selected the optimal $K$-value for the episode and each recall topic proportions matrix using the formula described in *Methods*. This computation resulted in a curve for each matrix, describing the Wasserstein distance between the distributions of within-event and across-event topic vector correlations, as a function of $K$.
